# Mitochondria isolated from male skeletal muscle contain a distinct population of miRNA that are differentially expressed following acute exercise

**DOI:** 10.1101/2024.10.13.617681

**Authors:** Jessica L. Silver, Stella Loke, Danielle Hiam, Larry Croft, Megan Soria, Søren Nielsen, Séverine Lamon, Glenn D. Wadley

**Author notes:** **Corresponding authors:** Dr Jessica Silver, Prof Glenn Wadley.

## Abstract

Initially thought to localise at the cytosol and nucleus only, emerging evidence indicates that miRNAs also localise within the mitochondria where they could regulate diverse pathological and physiological processes. Therefore, the aim of the current study was to use small RNA sequencing to profile and compare the entire population of miRNAs in human skeletal muscle of healthy males in whole-tissue and in isolated mitochondria at rest and in response to acute endurance exercise. Twelve healthy males (age 26 ± 4 years, mean ± SD) cycled for 60 min at 70% VO_2peak_ and muscle biopsies were collected at rest, immediately after and 3 h after exercise. The mitochondria were isolated by immunoprecipitation, further purified, then the resident RNA was sequenced to assess the mitochondrial transcriptome. Small RNA sequencing revealed that mitochondria isolated from male skeletal muscle tissue contain a small and distinct population of miRNAs. Of the approximately 110 mature miRNAs detected in skeletal muscle mitochondria at each time-point, the canonical myo-miRs miR-1, miR-133 and miR-206 families constituted on average 45% of total mitochondria miRNA reads. However, none of these canonical myo-miRs were differentially expressed in mitochondria following endurance exercise. One miRNA, hsa-miR-146b-3p, was differentially expressed in both whole muscle tissue and mitochondria when adjusted for multiple testing (FDR <0.05). Future research is now required to investigate miRNA-mRNA interactions in the mitochondria of skeletal muscle tissue.

**KEY POINTS SUMMARY:**

- Emerging evidence suggests microRNA are localised in the mitochondria of skeletal muscle cells and may play a role in regulating mitochondrial function.
- We recently optimised an approach to isolate RNA from mitochondria of human skeletal muscle that is free from contaminating cytosolic RNA and suitable for RNA sequencing.
- In this study we examined the microRNA population from male skeletal muscle mitochondria before and in the hours following 60 minutes of moderate intensity cycling exercise.
- We detected around 110 microRNAs in skeletal muscle mitochondria, with the muscle enriched myo-miR such as miR-1, miR-133 and miR-206 families constituting almost half of the reads. However, only one microRNA, hsa-miR-146b-3p, was differentially expressed whereby it increased ∼ 10-fold following exercise.
- The results provide new knowledge into how mitochondria might be regulated at the subcellular level and in response to physiological stressors such as exercise.

## INTRODUCTION

Skeletal muscle is a dynamic tissue that is characterised by high mitochondrial content (1). The mitochondria retain their own genetic code (2, 3) and are populated by over 2,000 nuclear-encoded proteins that are altogether required for optimal mitochondrial function. Endurance training results in adaptations that are beneficial to muscle tissue via improvements in mitochondrial content and function (4). In contrast, mitochondrial dysfunction is associated with chronic metabolic conditions such as cardiovascular disease and type 2 diabetes (4).

The metabolic stress associated with endurance exercise training promotes a shift towards a more oxidative phenotype in skeletal muscle that is largely due to an increase in mitochondrial biogenesis (4, 5). Although this increase and the subsequent increases in mitochondrial content and activity are associated with sustained endurance exercise training, over 200 skeletal muscle genes are up regulated in the 48-h following a single bout of endurance exercise (6). However, exercise-induced changes in gene expression at the whole skeletal muscle level only tell part of the picture. For example, Ppargc-1a muscle mRNA levels increase by 10-fold 3h following acute endurance exercise (7–9) whilst its protein levels can remain unchanged (8) or perhaps increase by up to 50% (9, 10). During this same period, the PPARGC-1 protein translocates to the nucleus, increasing its nuclear abundance 2-fold to exert its functions in the early recovery phase following acute endurance exercise (10). This suggests that the subcellular response to endurance exercise may be equally important as the whole cell response.

Non-coding RNAs such as micro RNAs (miRNAs) can fine-tune the regulation of transcript and protein abundance in mammalian cells and tissues (11). MiRNAs are short, non-coding RNA molecules that selectively recognise untranslated regions of a target mRNA (12) and most commonly repress translation by degrading a target mRNA (13). Within skeletal muscle, miR-1, miR-133a/b, miR-206 and miR-486 are highly abundant (14) and have various roles in the regulation of myocyte development, growth, regeneration and metabolism (12). MiR-1, miR-133a and miR-133b (15, 16), among others (17), are regulated in skeletal muscle following acute exercise. Loss-of-function models demonstrate that miR-133a knockdown reduces Ppargc-1α and Tfam translation (18), both of which are key regulators of the mitochondria biogenesis pathway. In addition, miR-1 can directly target and increase the expression of numerous mitochondrial-encoded transcripts (19, 20) whilst at the same time miR-1 is observed to repress nuclear encoded transcripts (20). This suggests that some ncRNAs can have opposing functions on transcripts depending on whether they are localised in the cytosol or mitochondria. The functional role of these miRNAs in relation to exercise-induced increased in mitochondrial content and activity is, however, still not well understood.

All known human miRNAs are first transcribed from the nuclear genome (21) before mature miRNAs are incorporated into RNA-induced silencing complexes (RISCs), usually within the cytoplasm (22). The localisation of miRNAs in subcellular compartments such as the mitochondria is increasingly well characterised in the heart (19, 20, 23, 24), liver (25) and kidneys (26, 27). While miRNA-mediated regulation of mitochondrial function and mitochondrial-encoded transcript abundance is an intriguing area of mitochondria physiology, there is currently no evidence detailing how miRNAs respond to metabolic stressors that induce a beneficial adaptation such as endurance exercise in skeletal muscle mitochondria. To date, evidence reporting the localisation of miRNAs in skeletal muscle mitochondria is limited. One group visually demonstrated by way of fluorescent in situ hybridisation that miR-365 is expressed in the mitochondria of primary human myoblasts in cell culture and profiled an additional 46 miRNAs in these mitochondria by micro array (28). Further research using high-throughput RNA sequencing techniques is however required to profile the entire population of miRNAs that are expressed in mitochondria isolated from human skeletal muscle tissue.

Therefore, the aim of the current study was to use small RNA sequencing to profile the entire population of miRNAs in human skeletal muscle of healthy males in isolated mitochondria and whole-tissue, and to investigate if miRNAs were differentially expressed at the whole-tissue or sub-cellular level in response to acute endurance exercise. A secondary aim was to use bioinformatics target prediction and pathway enrichment analyses to investigate putative associations between differentially expressed mitochondria-localised miRNAs and target transcripts and proteins regulating mitochondrial function in human skeletal muscle.

## METHODS

### Participant Recruitment

This study was approved by Deakin University Human Research Ethics Committee (EC 2021-223) and conforms to the *Declaration of Helsinki*. Written, informed consent was obtained from all participants before commencing exercise trials and sampling procedures. Twelve healthy males (age 25.6 ± 3.6 years; body mass index 25.5 ± 3.7 kg.m^-2^; VO_2peak_ 42.2 ± 6.1 ml.kg^-1^.min^-1^; mean ± SD) were recruited. Participants were excluded if they were smokers, had a family history (parent or grandparent) of heart disease, a bleeding disorder, BMI <18 or >30 kg.m^-2^; a peak oxygen consumption (VO2peak) of >55 ml.kg^-1^.min^-1^ or a pre-existing musculoskeletal injury or respiratory condition that could impact their ability to cycle at both maximal and sub-maximal intensities.

### VO_2_peak and Exercise Testing

Participants reported to the laboratory on two occasions. In the first visit, participants reported to the laboratory, having performed no moderate-vigorous exercise in the preceding 48 h and underwent a maximal incremental exercise test on an electronically braked cycle-ergometer *(Lode, Groningen, NED)* to determine VO_2_peak. Participants cycled at a self-selected cadence throughout the test. Cycling began at 75 W and increased by 50 W every three minutes for nine minutes. Thereafter, the workload increased by 25 W every minute until volitional fatigue. Individual VO_2_peak was calculated using data from expired gases using an Innocur breath-by-breath metabolic system (Innovision, Glammsbjerg, DEN). VO_2_peak was defined as the highest VO_2_ recorded. At least one week later, participants reported to the laboratory at 7:00am following an overnight fast and performed a 60-minute bout of cycling at 70% VO_2_peak.

### Skeletal Muscle Tissue Collection

Skeletal muscle samples were obtained via muscle biopsy using the percutaneous muscle biopsy technique with a Bergstrom needle, modified to include suction (29, 30). Briefly, the skin was anaesthetised with 1% Xylocaine, and incisions were made through the skin and muscle fascia. Three muscle biopsies were obtained from the vastus lateralis of the same thigh, from three separate incisions approximately 20 mm apart. Skeletal muscle biopsies were taken at rest (pre), immediately after exercise at 70% VO_2_peak (post), and following 3 h of passive recovery (3h post). Two approximately 60 mg portions of freshly obtained skeletal muscle were blotted free of blood and immediately processed for the isolation of mitochondria. The remaining skeletal muscle was immediately frozen in liquid nitrogen and stored in liquid nitrogen for future analyses.

### Mitochondria Isolation

Each 60 mg portion of freshly obtained skeletal muscle was suspended in 1 mL ice-cold Lysis Buffer (Mitochondria Isolation Kit, human, 130-094-532, Miltenyi Biotec, CA, USA) containing 5 μL/mL Protease Inhibitor Cocktail (P8340, Sigma-Aldrich, MA, USA) and homogenised on ice for 10 strokes (up and down) using a Dounce homogeniser and mechanical rotor at low speed. Mitochondria were isolated using the MACS Mitochondria Isolation Kit (human tissue) (130-094-532, Miltenyi Biotec, CA, USA). Briefly, the lysate was incubated with 50 μL ice-cold magnetic microbeads conjugated to anti-TOM22, a protein of the outer mitochondrial membrane. This immunoprecipitation approach isolates mitochondria free from golgi/peroxisome protein complexes (31) that may contain cytosolic RNA. Following isolation, the mitochondrial fraction was resuspended in 50 μL ice-cold Storage Buffer (130-094-532, Miltenyi Biotec, CA, USA) and incubated with 312 μg/mL RNase-A (19101, Qiagen, Hilden, Germany) for 1 h at 37°C. Following incubation, RNase-A was degraded by the addition of proteinase-K (3 mAU/mL, 19131, Qiagen, Hilden, Germany) and inverting the tube briefly. The solution was centrifuged at 8,000*g* for 10-minutes at 4°C. The supernatant was aspirated, and the mitochondria pellet was washed twice with 100 μL ice-cold Storage Buffer (130-094-532, Miltenyi Biotec, CA, USA), and centrifuged at 13,000*g* for 2-minutes at 4°C between washes. The mitochondria pellet was resuspended in 100 μL storage buffer (130-094-532, Miltenyi Biotec, CA, USA) and stored at -80°C until further analyses. We have previously developed this method for obtaining a highly pure fraction of mitochondria from skeletal muscle tissue and optimised this approach for high throughput transcriptomic sequencing on this subcellular fraction (32). The incubation of isolated mitochondria with RNase-A digests any contaminating extra-mitochondrial RNA to ensure that only RNA species contained within the mitochondria membranes are identified in downstream analyses (32).

### RNA Extraction

For RNA extraction from isolated mitochondria, frozen samples were suspended in 5x volumes of TriReagent (R2060, Zymo, CA, USA), thawed on ice and then centrifuged at 16,000x*g* for 2 min at 4°C to pellet the mitochondria. The mitochondria pellet was first sheared 20 times through a fine pipette tip. Total RNA was extracted from isolated mitochondria using the Zymo Direct-Zol Micro Prep kit (R2060, Zymo, CA, USA) with on-column DNase-I digest (E1010, Zymo, CA, USA) as per the manufacturer’s instructions. Mitochondrial RNA was eluted in 12 μL nuclease-free water and quantified by gel electrophoresis using the TapeStation High Sensitivity RNA screentape and reagents (5067-5579 and 5067-5580, Agilent Technologies, CA, USA). Mitochondria total RNA concentrations ranged between 0.5-8.6 ng/µL (mean ± SD, 2.3 ± 2.1 ng/µL). A representative electropherogram of total RNA extracted from human skeletal muscle mitochondria is presented in Supporting Information-Figure 1A.

For RNA extraction from skeletal muscle tissue, 10-15 mg frozen human skeletal muscle tissue was submerged in 400 μL ice-cold 1x DNA/RNA protection reagent (T2010, New England Biolabs, MA, USA) containing 700-800 mg 1.0mm zirconia/silica beads (11079110z, BioSpec Products, St. Helena, Australia). The tissue was homogenised with mechanical disruption at 6,500rpm for 30 s (MagnaLyser, Roche Diagnostics, NSW, Australia). Next, 30 μL Prot K Reaction Buffer (T2010, New England Biolabs, MA, USA) plus 15 μL Prot K (T2010, New England Biolabs, MA, USA) was added to the tissue homogenate and incubated at 55°C for 5 min. Following incubation, the lysate was briefly vortexed and then centrifuged at 16,000x*g* for 2 min. The supernatant was transferred to a new tube. RNA extraction was then performed using the Monarch Total RNA Mini Prep Kit (T2010, New England Biolabs, MA, USA) with on-column DNase-I digest as per the manufacturer’s instructions. RNA was eluted from the column in 30 μL nuclease-free water. The eluted RNA was tested for purity and concentration on a Nanodrop One (Thermo Fisher Scientific, MA, USA). If required, RNA was diluted below 25 ng/μL and then the RNA integrity number (RIN) was determined using the HS RNA screentape and reagents (5067-5579 and 5067-5580, Agilent Technologies, CA, USA) on the Agilent 4200 TapeStation.

In RNA from whole tissue, A260/280 and A260/230 were ≥2.0 and ≥1.8, respectively, and all RINs were ≥7.0 (mean ± SD, 7.4 ± 0.2). The skeletal muscle total RNA concentrations ranged between 22.4-76.2 ng/µL (mean ±SD, 49.8 ± 152.4 ng/µL). A representative electropherogram of total RNA extracted from human skeletal muscle tissue is presented in Supporting Information-Figure 1B.

### Small RNA Sequencing

#### Small RNA Library Preparation (Isolated Mitochondria)

Small RNA libraries were prepared using the NEBNext Multiplex Small RNA Library Prep Set (E7560S, New England Biolabs, MA, USA) from 13.4 ± 11.2 ng total mitochondrial RNA (range for *n*=3 42.9-51.6 ng when compared to 9.8 ng on average in all other samples) as we have described previously (32). Briefly, the 3’ and 5’ adapters and reverse transcription primer were diluted 0.3x, and 20 end-point PCR cycles were conducted. Following PCR amplification, the cDNA library was cleaned using the Monarch PCR & DNA Cleanup Kit (5 μg) (T1030L, New England Biolabs, MA, USA) using the 7:1 ratio of binding buffer:sample as per the manufacturer’s instructions and was eluted in 27.5 μL nuclease-free water. The eluate containing the purified cDNA construct was first quantified using the 1X dsDNA HS Assay Kit (Q33231, Thermo Fisher Scientific, CA, USA) on a Qubit 4.0 Fluorometer. Fragment size distribution was then assessed on the Agilent 4200 TapeStation using the HS-D1000 screentape and reagents (5067-5584 and 5067-5585, Agilent Technologies, CA, USA). All mitochondrial libraries contained the anticipated peak corresponding to adapter-ligated miRNAs (approximately 160 bp, Supporting Information-Figure 2A), which accounted for 32.8 ± 13.8% of the total cDNA library. A total of 33 libraries (pre-exercise, n=9; post-exercise, n=12; 3h post-exercise, n=12) passed quality checks.

#### Small RNA Library Preparation (Whole-tissue)

Small RNA libraries were also prepared from 299 ± 167 ng (n=35) total RNA extracted from whole skeletal muscle tissue using the NEBNext Multiplex Small RNA Library Prep Set (cat# #E7560S, New England Biolabs.) with 3’ and 5’ adapters and reverse transcription primer diluted 0.5x, and 15 PCR cycles to amplify the adapter-ligated library. The cDNA was cleaned, quantified and assessed as described above for mitochondrial small RNA libraries. All whole skeletal muscle libraries contained the anticipated peak corresponding to adapter-ligated miRNAs (approximately 160 bp, Supporting Information-Figure 2B)

#### Size Selection

In our hands, purification of adapter-ligated miRNAs via gel-excision has minimised the presence of adapter-dimer (32). However, these samples previously required re-amplification, bead-based clean-up and additional gel-excision to produce sufficient amounts of library required for sequencing on the NovaSeq6000 platform.

Small RNA libraries were first placed into one of two equimolar library pools (corresponding to isolated mitochondria and whole tissue libraries, respectively). To maximise the amount of library available for sequencing without the need for re-amplification and additional clean-up of the gel-excised fragments, aliquots of the pooled mitochondrial libraries (5×27.5 μL aliquots) and pooled skeletal muscle libraries (3×27.5µL aliquots) were run in separate lanes of a 6% Novex TBE-polyacrylamide gel (EC6265BOX, Thermo Fisher Scientific, MA, USA), with 5 μL Quick-Load pBR322 DNA-MspI Molecular Marker (E7560S, New England Biolabs, MA, USA) run in a separate lane. The gel was then incubated with 50 mL 1X TBE spiked with 1X SYBR Gold Nucleic Acid Gel Stain (S11494, Thermo Fisher Scientific, MA, USA) for 10-minutes at room temperature with gentle shaking. DNA bands corresponding to the 140 (adapter-ligated miRNAs) and 190 (other small RNAs) bp markers were manually excised under exposure to UV-Blue (365 nm) light, suspended in 250 μL gel elution buffer (E7560S, New England Biolabs, MA, USA) and incubated for at least 4 h on a rotating wheel at room temperature. The longer incubation time than indicated in the manufacturer’s protocol (2 h) purportedly increases elution of small fragments of DNA (up to 500 bp) from the gel (33). Following incubation, cDNA was precipitated overnight at -20°C (33) and then the solution was centrifuged at 16,000xg for 60 min to pellet the cDNA fragment. After two sequential washes of the cDNA pellet in freshly prepared 80% ethanol, the gel-excised regions were resuspended in 12 µL TE buffer. Fragment size distribution was then assessed on the Agilent 4200 TapeStation using the HS-D1000 screentape and reagents (5067-5584 and 5067-5585, Agilent Technologies, CA, USA). After quantification and quality checks of the gel-excised regions, each aliquot was combined into pools representing the mitochondrial 140 and 190 bp and whole muscle 140 and 190 bp gel-excised regions.

#### Small RNA Pre-Sequencing (MiSeq) and Data Pre-Processing

First, RNA pre-sequencing was performed by Charles River Laboratories (VIC, Australia). The pre-sequencing run was used for validation purposes to confirm that i) all unique indexes efficiently ligated to mitochondrial RNA samples during library preparation; ii) assess sample distribution; and iii) confirm that residual adapter-dimer did not limit the detection of miRNAs within the pool. First, a 5 pM loading pool was prepared containing 25% gel-excised miRNA region (mitochondria pool only), 25% 190 bp region (mitochondria pool only) and 50% Phi-X spike-in and was pre-sequenced on a MiSeq Reagent Nano Kit v2 (MS-102-2001 Illumina, SD, USA) single-end 1×51 bp run. Following pre-sequencing, quality control and pre-processing of fastq files (read1 only) were performed using fastp (fastp v0.20.0) (34) to remove the read1 adapter sequence (AGATCGGAAGAGCACACGTCTGAACTCCAGTCA) and filter poor quality reads. Next, reads were mapped using blast (v2.9.0+) with switch -task blastn and word length set to 19 nt to miRbase human mature miRNAs (mature.fa; accessible from miRbase (v22.1) (2); downloaded March 12, 2018). Raw read counts for each mature miRNA were counted using a short perl script.

Within each library, the total number of sequenced reads was not significantly different in response to acute endurance exercise (*p*<0.69; Supporting Information-Figure 3A). The proportion of reads mapping mature human miRNAs (mean ± SD, 3.76 ± 8.25%; range, 0.24-27.65%, Supporting Information-Figure 3B) was not significantly different in response to acute exercise. For five small RNA libraries, the proportion of reads mapping miRNAs was between 20-30% and was well above the range presented for most samples (approximately 0.6%, on average; Supporting Information-Figure 3B). Despite these clear outliers, the total number of sequenced reads was consistent with all other samples suggesting that there were no differences in the sequencing depth across all libraries. Therefore, no modifications to the library pools were made before proceeding to the main RNA sequencing on the NovaSeq platform.

#### Small RNA Sequencing (NovaSeq 6000) and Data Pre-Processing

Human skeletal muscle mitochondria and human whole skeletal muscle tissue pools were subsequently loaded on separate lanes on a SP flow cell on the NovaSeq 6000 XP (v1.5) platform. A 2nM loading pool was prepared for each of the whole tissue and isolated mitochondria samples, respectively. Each pool contained 50% gel-excised miRNA region, 25% gel-excised 190 bp region and 25% Phi-X spike-in and was sequenced on a 2×51bp run. Following sequencing, quality control and data pre-processing were performed for read1 only as described for pre-sequencing above.

A small RNA quality score was generated for each library and is presented the percentage of reads mapping exons (coding sequences) per 10,000 reads. One-way, repeated measures ANOVAs with statistical significance set at 0.05 were used to identify significant differences in total sequenced reads, the proportion of reads mapping coding sequences, the proportion of reads mapping mature miRNAs and the total number of miRNAs over time. RNA-sequencing datasets have been deposited at the NCBI Gene Expression Omnibus for the small RNA sequencing for whole skeletal muscle (accession number GSE277549) and for isolated mitochondria (accession number GSE220280).

#### Total RNA Sequencing (Whole Skeletal Muscle Tissue)

Total RNA extracted from whole skeletal muscle tissue was prepared for total RNA sequencing externally (Macrogen, South Korea) and served two purposes in the current study: 1) to investigate if the exercise protocol was sufficient to induce the metabolic stress and increase expression levels of transcripts known to be upregulated following an acute bout of endurance exercise, and 2) to profile all transcripts that are expressed in skeletal muscle tissue pre-, post- and 3h post-exercise for correlation analyses.

Total RNA libraries depleted of ribosomal RNA were prepared using the TruSeq stranded Total RNA Library Preparation with RiboZero (human/mouse/rat). Total RNA libraries were placed in an equimolar pool and sequenced on the NovaSeq 6000 (v1.5) platform on a 150 bp paired-end run. Following sequencing, sequence quality was inspected using fastqc (v0.11.9, Babraham Bioinformatics, UK) and then tabulated using multiqc (v1.13, (35)). RNA-seq reads were mapped to the human transcriptome (GRCh38.p13, downloaded from ENSEMBL v38) using kallisto v0.46.1. Data were then read into RStudio (v4.2.1). Duplicate lane data were aggregated, and transcript level counts were aggregated to gene level counts based on ENSEMBL ID. Genes with an average of less than 10 reads across samples were excluded from downstream analysis. RNA-sequencing datasets from this study have been deposited at the NCBI Gene Expression Omnibus for the whole-transcriptome in whole skeletal muscle (GSE276889).

### RNA Sequencing Data Analysis

#### Differential Expression Analysis

Analysis of differential miRNA expression in mitochondria and whole-tissue samples, and mRNA expression in whole-tissue samples only, obtained pre-, immediately post- and 3h post-exercise was performed using DEseq2 (v1.36) in Rstudio (v4.2.1). DESeq2 uses raw, non-normalised read counts as its input and integrates a backend normalisation that calculates the geometric mean for each gene across all samples and internally corrects for library size prior to differential expression analysis (36). Pre-filtering was undertaken to remove genes or miRNAs with less than 10 normalised read counts across all samples. The false discovery rate (FDR, adjusted for multiple testing using the Benjamini and Hochberg approach) and log_2_ fold-change (relative to pre- or immediately post-exercise) are presented alongside the raw p-value for each miRNA (37). MiRNAs and mRNAs with an FDR<0.05 and log_2_FC >1 or <-1 were identified as differentially expressed. Heatmaps to reveal patterns of gene expression were generated using the “ComplexHeatmap” package in R (v2.18.0) (38). Scripts used for differential expression analysis and data visualisation can be accessed from https://github.com/jessicasilver/miRNAs. Data was visualised in GraphPad Prism v8.0.0.

#### MRNA and miRNA Correlation Analysis

First, the differential expression analysis results for mRNA expression in whole-tissue across the two contrasts (3h post- vs pre-exercise) were used for the correlation analysis. From these results, the fold-change, raw p-value and ensemble gene IDs (n=15,198) were extracted. Next, fold-changes, p-values and ensemble gene IDs were extracted from the differential expression analysis results of skeletal muscle and mitochondria miRNAs, respectively, across the same two time-points. 772 and 387 gene IDs were extracted for skeletal muscle and mitochondrial samples, respectively. Finally, the miRanda target database for homo sapiens (hg19) was supplied by the mirTarRNASeq package (v1.8.0) (39) and used for miRNA-mRNA target identification and interaction analysis. The miRanda algorithm finds miRNA targets in two phases. Phase one includes a dynamic programming local alignment between the miRNA and the reference sequence that is scored based on the sequence complementarity. Phase two then takes the high-scoring alignments from phase one as input, estimates the thermodynamic stability of RNA duplexes and then performs a statistical analysis of the folding behaviour of the fictional single-stranded RNA from the query sequence (see (40) for detailed explanation of the algorithm). The significant gene targets were used to perform a hypergeometric test to determine significantly overrepresented pathways within the GO Biological Pathways gene set from the Molecular Signatures Database curated by the Broad Institute. This gene set was extracted using the “msigdbr” package in R (v7.5.1) (41).

#### Time-Course Correlation Analysis Using miRTarRNASeq

The time-course correlation analysis was performed using the mirTarRnaSeq R package (version 1.8.0) and was done separately for the skeletal whole-muscle mRNA to whole-muscle miRNA pair and for the skeletal whole-muscle mRNA to mitochondrial miRNA pair. This correlation analysis takes the log2 fold changes of the DE analysis results and performs a Pearson correlation across the inputs. After a threshold of p-value < 0.05 from the original DE analysis result table was applied, 2752 gene IDs passed for the muscle mRNAs from the original 15,198 and was used as input.

## RESULTS

### Participant Exercise Performance

All participants completed the full 60-minute exercise bout and maintained a mean (± SD) power output of 127 ± 21 watts that corresponded to 70.6 ± 4.1% of individual VO_2_peak during exercise.

### Whole-Muscle Transcriptomic Response to Acute Endurance Exercise

We first used RNA-Seq to describe the whole-muscle transcriptomic response to an acute endurance exercise bout. Of over 15,000 genes detected in skeletal muscle, 412 and 208 were up-regulated (FDR<0.05) 3h post-exercise when compared to pre-exercise and immediately post-exercise, respectively. The top 70 differentially expressed mRNAs in response to the acute exercise bout (FDR <0.05, |log_2_FC| >1) are shown in Figure 1. *PPARGC-1α* expression displayed the highest adjusted p-value of all differentially expressed genes and increased by approximately 6-fold 3h post-exercise when compared to pre- and immediately post-exercise (FDR <0.0001), consistent in magnitude to prior studies when assessed using qPCR (7, 8, 47, 48). This confirms that the exercise protocol was sufficient to induce the metabolic stress associated with acute endurance exercise.

**Figure 1.**
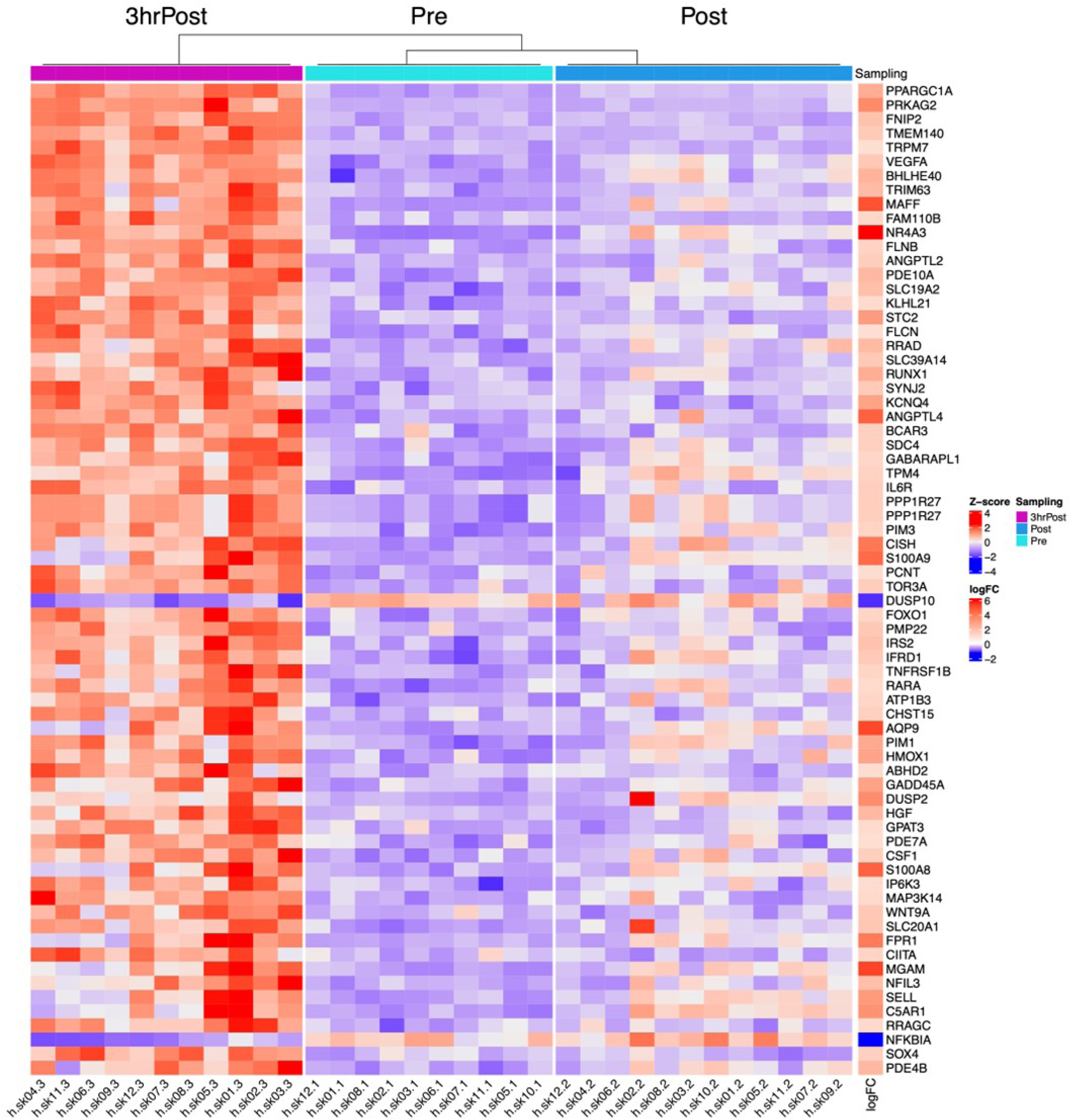
Differential mRNA expression in skeletal muscle tissue in response to 60 minutes of cycling at 70% VO_2_peak. Heatmap depicts z-scores for the top 70 significant mRNAs following differential expression analysis (padj<0.05, L2FC>1 or <-1) ordered by increasing adjusted p-values and generated using the 3h post- vs pre-exercise contrast. The samples were grouped by time point and clustered using Euclidean distance.

### MiRNA Expression in Skeletal Muscle Mitochondria and Whole Skeletal Muscle Tissue

In small RNA libraries prepared from skeletal muscle mitochondria collected pre-, post- or 3h post-exercise, there were no significant differences in the total number of reads (*p*=0.41; Figure 2A), the proportion of reads mapping transcriptome (5.41±1.64%, mean ± SD for all participants, *p*=0.41 over time; Figure 2B) or the proportion of reads mapping mature miRNAs (*p*=0.20; Figure 2C). Five unique small RNA libraries (from *n*=2 participants) showed a substantially higher proportion of reads mapping miRNAs (range=15.3-48.7% when compared to 0.8% on average in all other samples; Figure 2C). When visualised using PCA, these five samples were clear outliers (Figure 2D) and were subsequently excluded from all further analyses.

**Figure 2.**
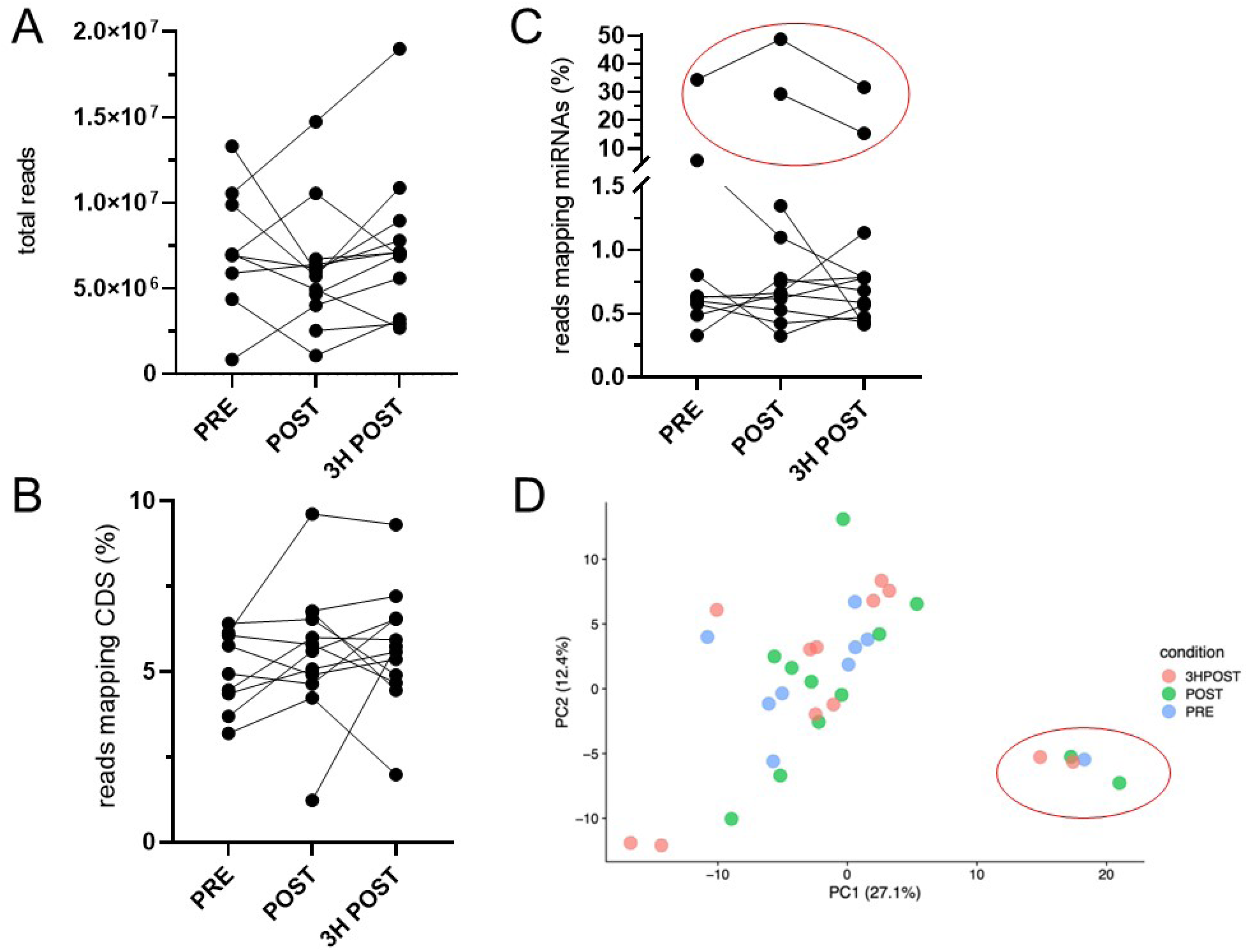
The A) total sequenced reads (p=0.41), B) proportion of reads mapping human-derived coding sequences (transcriptome, p=0.41), and C) proportion of reads mapping mature human miRNAs (p=0.20) were not significantly different between small RNA libraries prepared from skeletal muscle mitochondria pre-, post- or 3h post-exercise. One-way, repeated measures ANOVAs with p-value <0.05 were used to identify significant differences. Small RNA libraries for 5 small RNA libraries from n=2 participants (circled in red) contained a higher (approximately 30%) proportion of reads mapping miRNAs when compared to all other libraries (approximately 0.5%). D) PCA plot shows mitochondrial fraction variance from three time points, with PC1 and PC2 explaining 27.1% and 12.4% of the variance, respectively. Five samples in the lower right (circled in red), diverging significantly from others, were identified as outliers from all conditions and removed to maintain data integrity for downstream analysis.

At the time of analysis, 2,656 mature miRNAs were annotated to the human genome in miRbase (v22.1) (42). The skeletal muscle-enriched hsa-miR-1-3p was the most highly expressed species in both whole muscle and isolated mitochondrial fraction. Hsa-miR-1-3p was detected at levels approximately 7- and 25-fold higher than any other miRNA in the isolated mitochondria and whole skeletal muscle libraries, respectively (Figure 3A, B). Other skeletal muscle-enriched miRNAs such as hsa-miR-133a, hsa-miR-133b and hsa-miR-206 were among the top 25 most highly expressed miRNAs in both isolated mitochondria and whole skeletal muscle but displayed considerably lower expression levels when compared to hsa-miR-1-3p (Figure 3A, B). In whole skeletal muscle tissue, approximately 255 miRNAs were detected at each time-point. The total number of miRNAs was not significantly different in response to the acute endurance exercise bout (Figure 3D). In contrast, in isolated mitochondria only half of this number of miRNAs were detected at any of the time points (pre-exercise, 114 ± 5 post-exercise, 109 ± 6; 3h post-exercise, 115 ± 5 mean ± SD) and was significantly different over time (p=0.035, Figure 3C). However, multiple comparisons testing revealed no significant differences between individual timepoints (Figure 3C).

**Figure 3.**
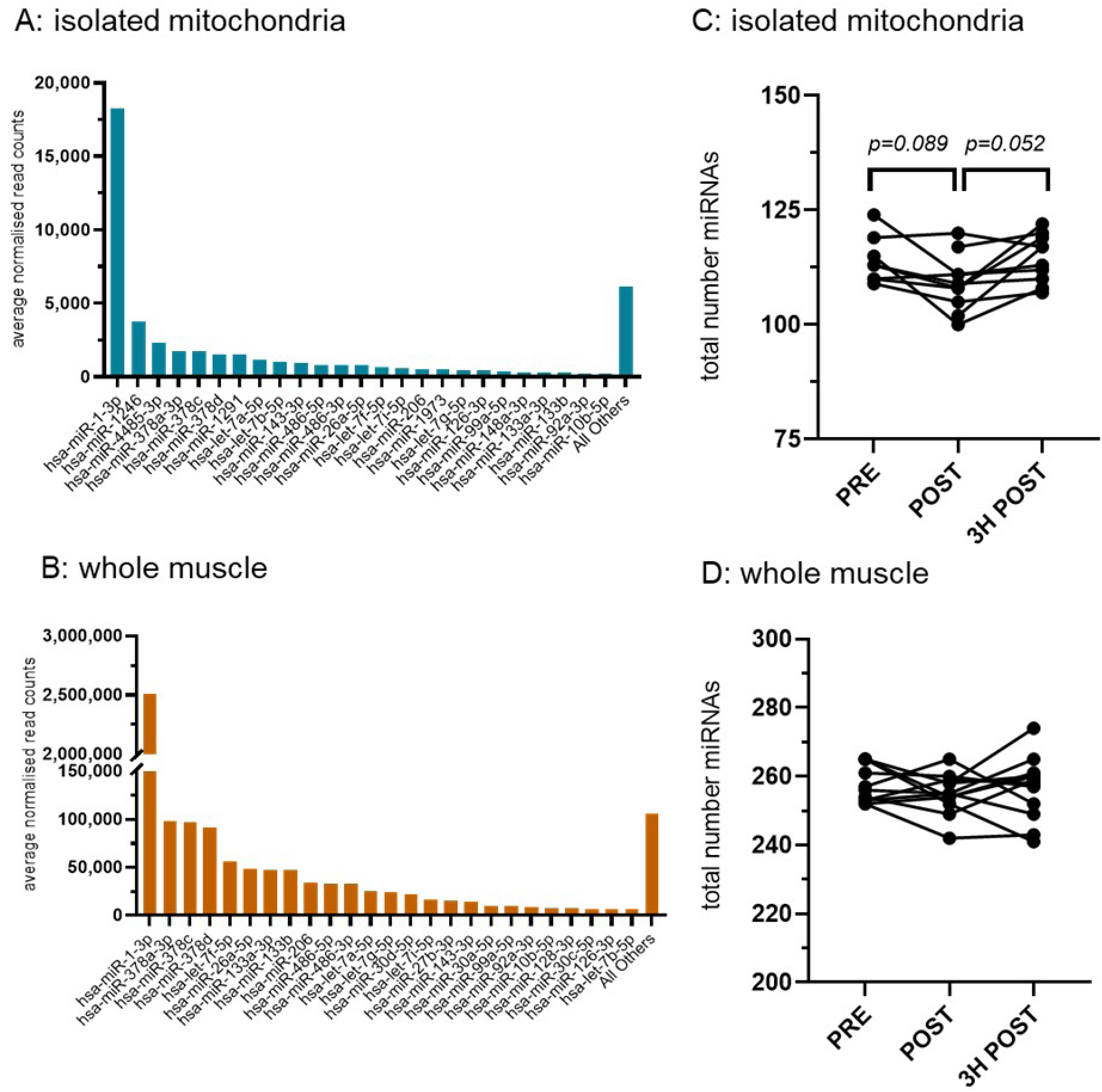
The top 25 most highly expressed miRNAs in A) isolated mitochondria and B) whole skeletal muscle tissue. C) The total number of individual miRNA species in mitochondria isolated from skeletal muscle tissue was significantly different over time (p=0.035), although Tukey’s multiple comparison tests did not reveal any significant differences between timepoints (p-values on graph). D) There were no significant differences in response to the exercise bout in skeletal muscle (p=0.588). One-way, repeated measures ANOVAs with p-value <0.05 were used to identify significant differences in the total number of miRNAs over time. Data displayed is the Trimmed Mean of M-values (TMM) normalised read count.

### Differential Expression of miRNAs in Whole Skeletal Muscle Tissue Following Acute Endurance Exercise

In whole skeletal muscle, a total of 70 (immediately post- vs pre-exercise), 39 (3h post- vs pre-exercise) and 14 (3h post- vs immediately post-exercise) miRNAs returned a raw p-value of less than 0.05 prior to FDR adjustment (Supporting Information Table 1). Of these, six mature miRNAs returned an adjusted p-value (FDR) of less than 0.05 for at least one of the contrasts. Hsa-miR-223-3p (Figure 4B) and hsa-miR-223-5p (Figure 4C) expression was significantly higher (FDR<0.05) both 3h post- and immediately post-exercise when compared to pre-exercise. Similarly, hsa-miR-15b-5p (Figure 4D) and hsa-miR-106b-3p (Figure 4E) expression were significantly higher (FDR<0.05) immediately post-exercise when compared to pre-exercise. Hsa-miR-146b-3p expression was significantly higher by 3h post-exercise when compared to immediately post-exercise (Figure 4F), while hsa-miR-452-5p was higher pre-exercise than at any other time point (Figure 4G). The canonical myomiRs (hsa-miR-1-3p, miR-133a, hsa-miR-133b and hsa-miR-206) were not differentially expressed at the whole skeletal muscle level in response to the acute exercise bout.

**Figure 4.**
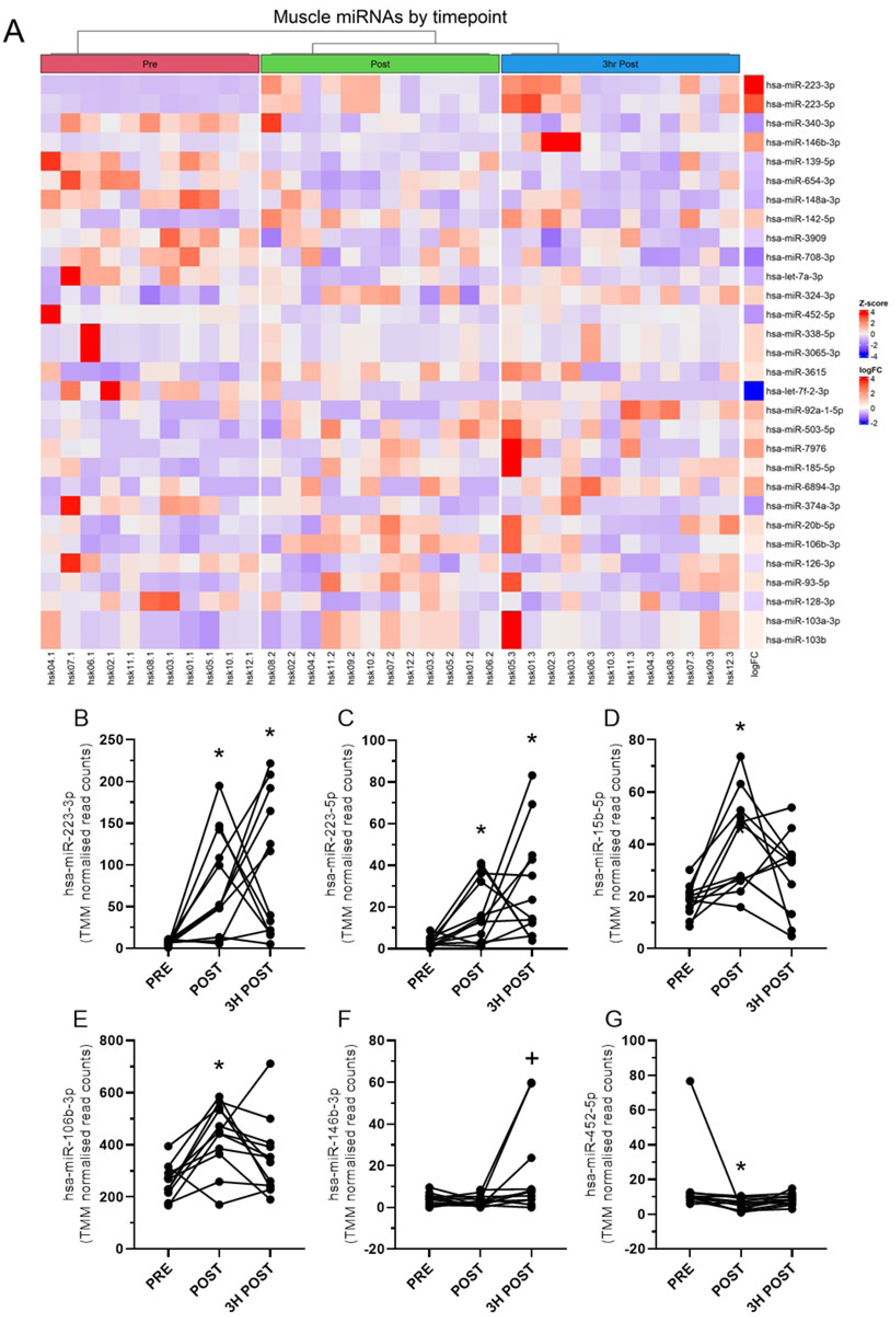
MiRNA expression in whole skeletal muscle tissue following 60 minutes of cycling at 70% VO_2_peak. A) Heatmap of miRNA expression in whole skeletal muscle tissue for miRNAs detected. The samples are grouped according to time points and are clustered using the Euclidean distance. B-G) Differentially expressed miRNA in whole skeletal muscle. Read counts were normalised using the Trimmed Mean of M-values (TMM) normalisation and analysed for differential expression using DESeq2. Individual TMM normalised read counts shown for each participant. * denotes FDR<0.05 versus PRE. + denotes FDR<0.05 versus POST.

### Differential Expression of miRNAs in Skeletal Muscle Mitochondria Following Acute Endurance Exercise

In isolated mitochondria, a total of 21 (immediately post- vs pre-exercise), 13 (3h post- vs pre-exercise) and 20 (3h post- vs immediately post-exercise) miRNAs returned a raw p-value of less than 0.05 prior to FDR adjustment (Supporting Information-Table 2). Eleven of these miRNAs returned a raw p-value of less than 0.05 prior to FDR adjustment for at least 2 contrasts (Figure 5B). Only one miRNA, miR-146-3p, returned a significant p-value for the immediately post to 3 hours-post contrast following FDR adjustment (Figure 5C). None of the canonical myomiRs such as hsa-miR-1-3p, hsa-miR-133a, hsa-miR-133b or hsa-miR-206 were differentially expressed in isolated mitochondria in response to acute endurance exercise.

**Figure 5.**
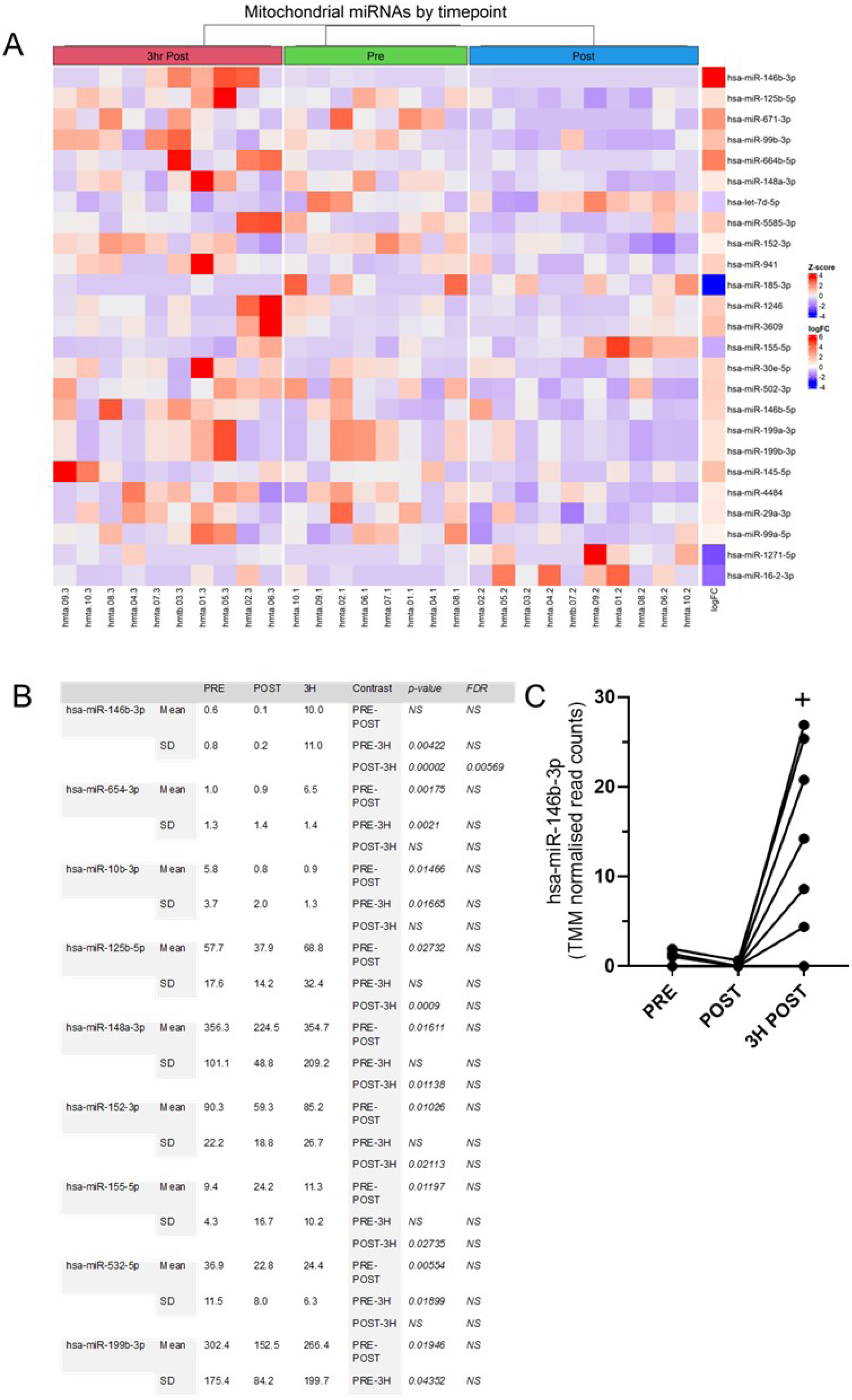
MiRNA expression in skeletal muscle mitochondria following 60 minutes of cycling at 70% VO_2_peak. A) Heatmap of mitochondrial miRNA expression for top 25 of 44 miRNAs detected ordered by p value and generated using the pre- vs 3h post-exercise contrast. Read counts were normalised using the Trimmed Mean of M-values (TMM) normalisation) and are grouped according to time points and are clustered using the Euclidean distance. B) Table of miRNAs with ≥ 2 contrasts with a raw p-value <0.05. C) Hsa-miR-146b-3p, the read counts were normalised using the TMM and analysed for differential expression using DESeq2. Individual TMM normalised read counts shown for each participant, + denotes FDR<0.05 versus POST.

### Correlation analysis or target prediction

Time course correlation analyses using miRTarRnaSeq were conducted across the two time points displaying the most differentially expressed mRNA profiles (Pre vs 3hrPost in Figure 1). Two correlation analyses were run independently. The whole muscle mRNA vs whole muscle miRNA correlation resulted in 105,207 significant interactions (p-value < 0.05) and 312 intersections with the miRanda database (Figure 6). An overrepresentation analysis was performed using the unique ensemble genes and resulted in 745 significantly enriched pathways (p-value < 0.05). The top 30 enriched GO biological pathways resulting from this overrepresentation analysis are displayed on Figure 7 and heavily feature pathways related to the immune and inflammatory response. The same treatment was applied on whole muscle mRNA and mitochondrial miRNA at the same time-points, which returned 52,386 significant mRNA-miRNA correlations and 236 intersections with the miRanda database (Figure 8) and 390 significant enriched pathways (p-value < 0.05) from the overrepresentation analysis. The top 30 enriched GO biological pathways resulting from this overrepresentation analysis display a similar overrepresentation of immune and inflammatory pathways (Figure 9).

**Figure 6.**
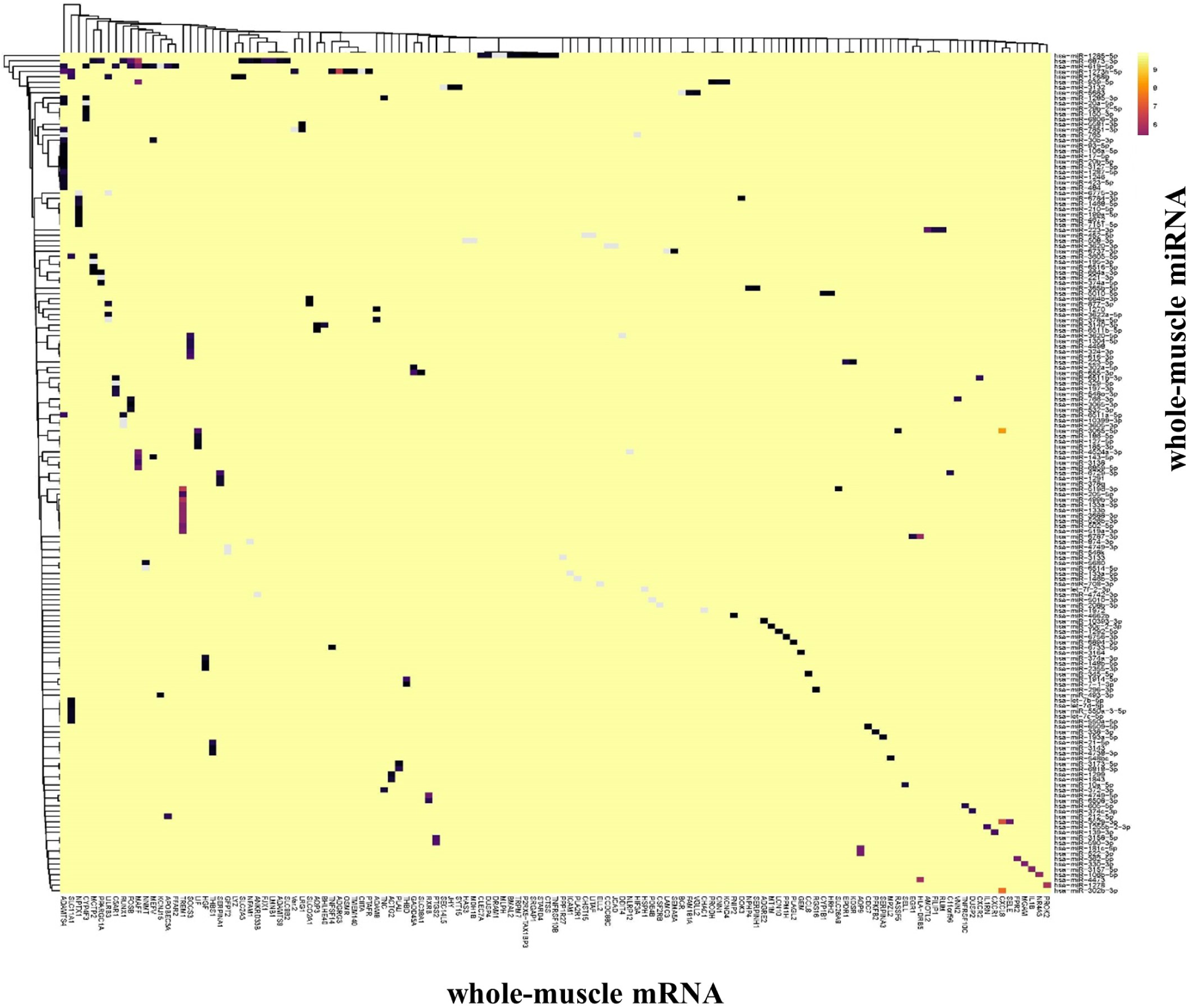
Heatmap of miRanda intersection with whole-muscle mRNA (x-axis) and miRNA (y-axis) correlations at 3 hours post- vs pre-exercise (p-value <0.05). The whole-muscle mRNA and miRNA correlation resulted to 105,207 significant interactions (p-value < 0.05) and 312 intersections with the miRanda database.

**Figure 7.**
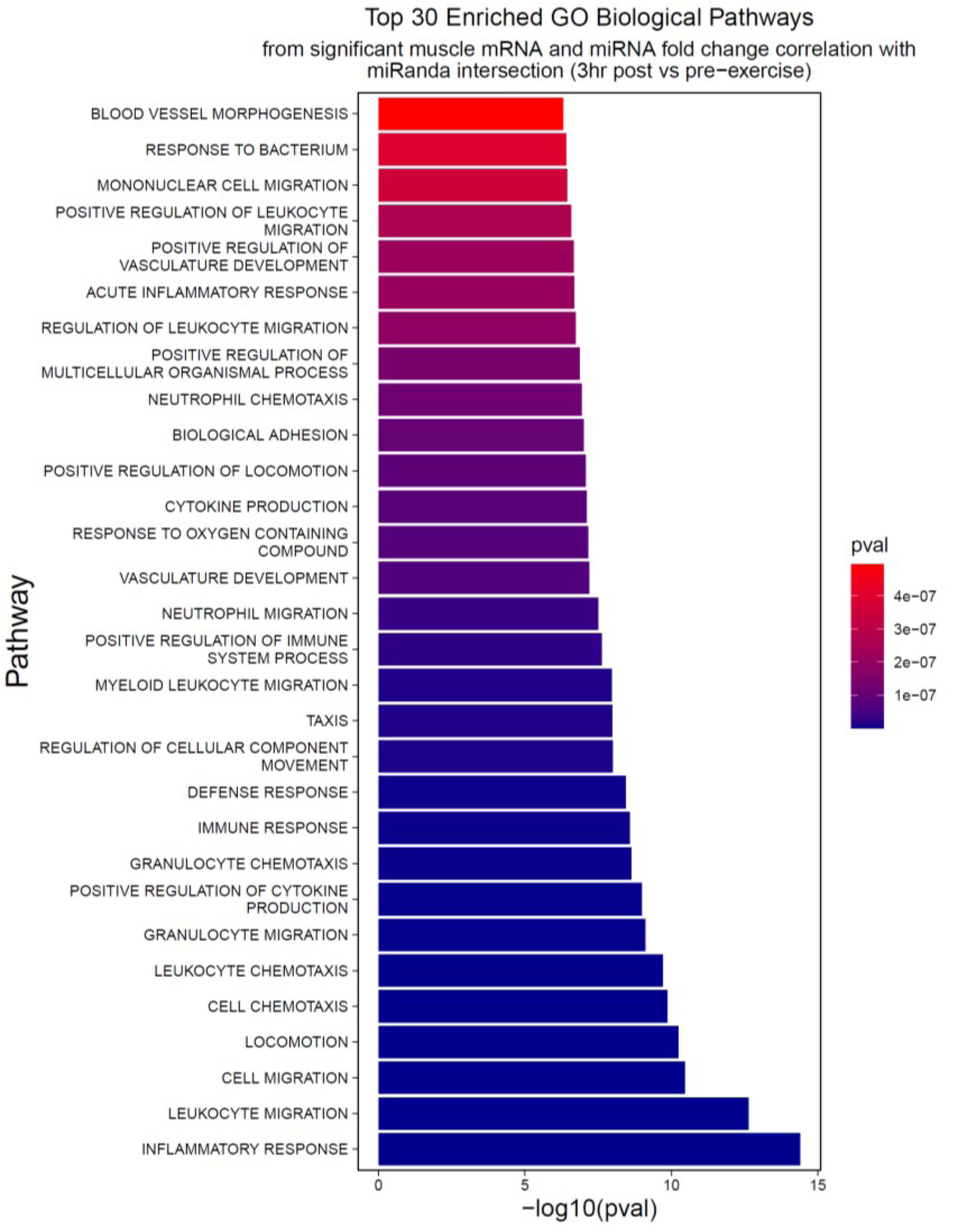
Top 30 enriched GO biological pathways from significant whole-muscle mRNA and miRNA fold-change correlation with miRanda intersection (3hr post vs pre-exercise).

**Figure 8.**
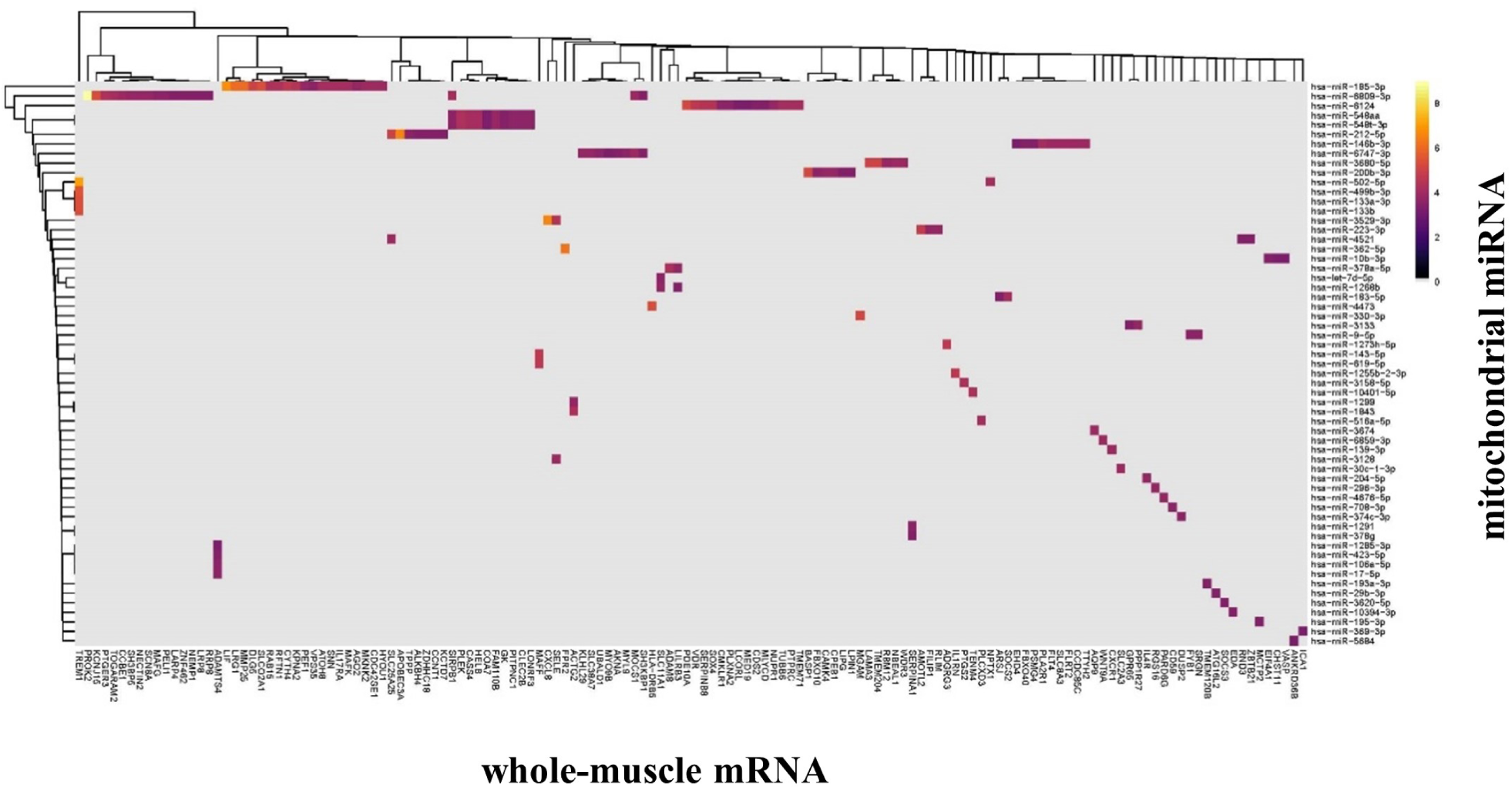
Heatmap of miRanda intersection with whole-muscle mRNA (x-axis) and mitochondrial miRNA (y-axis) correlations at 3 hours post- vs pre-exercise (p-value <0.05).

**Figure 9.**
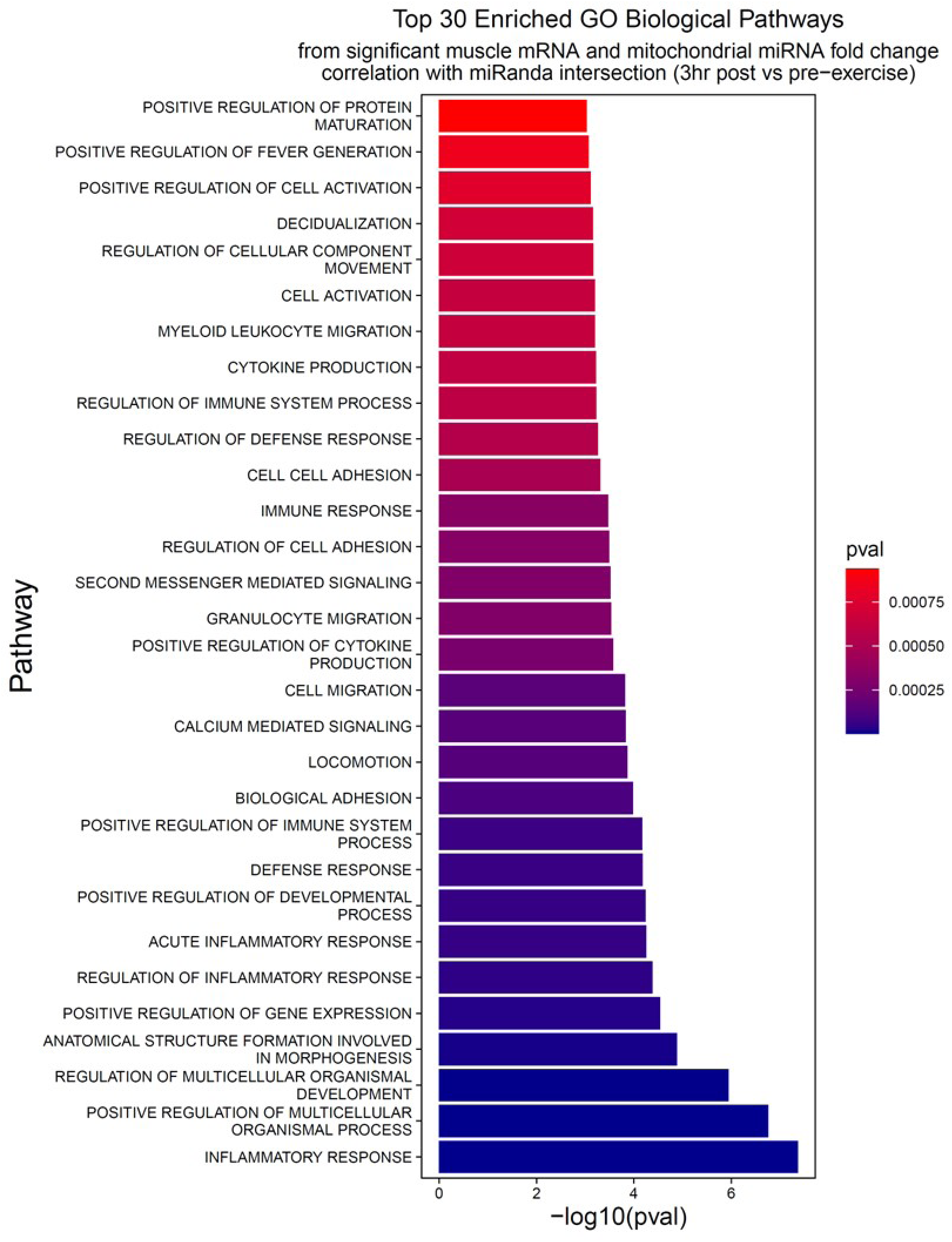
Top 30 enriched GO biological pathways from significant whole-muscle mRNA and mitochondrial miRNA fold-change correlation with miRanda intersection (3hr post vs pre-exercise).

Finally, the overrepresented pathways in both interactions were cross-checked and 221 significant (p-value < 0.05) enriched GO biological pathways were determined to be common to both, revealing a significant overlap between both (Figure 10).

**Figure 10.**
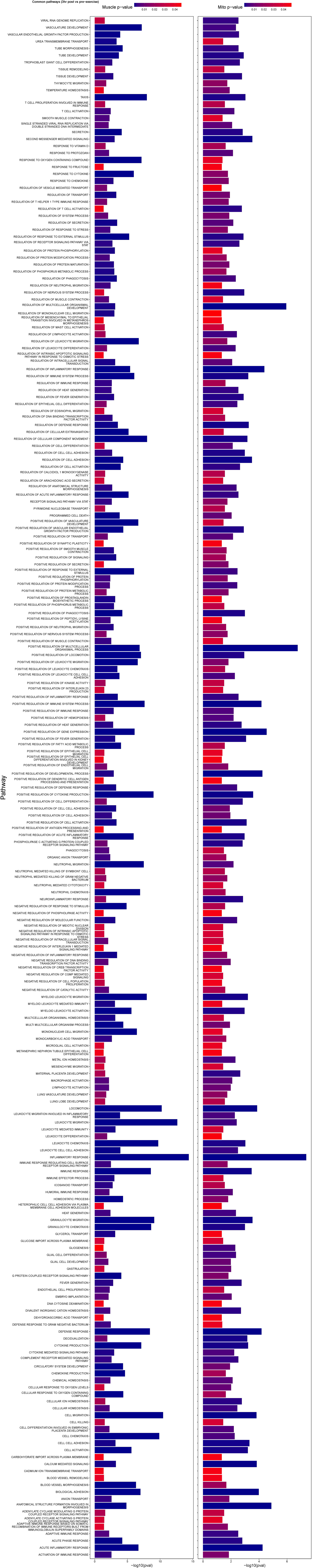
Common GO biological pathways between whole-muscle mRNA and mitochondrial miRNA fold-change correlations with miRanda intersection (3hr post vs pre-exercise).

## DISCUSSION

In the current study, small RNA sequencing revealed that mitochondria isolated from human skeletal muscle tissue contain a small and distinct population of miRNAs. Of the approximately 110 mature miRNAs detected in skeletal muscle mitochondria at each time-point, the canonical myo-miRs (miR-1, miR-133 and miR-206 families (43)) constituted on average 45% of total mitochondria miRNA reads. However, none of the canonical myo-miRs were differentially expressed in mitochondria following endurance exercise. Despite differential expression analysis identifying several miRNAs in isolated mitochondria that returned raw p-values less than 0.05 at multiple timepoints, only hsa-miR-146b-3p was differentially expressed when adjusted for multiple testing. Although experimentally validated interactions between miRNAs and mRNAs encoded by the mitochondrial genome are limited in the literature, the putative miRNA-mRNA associations found to be overrepresented following exercise were pathways relating to immune and inflammatory responses.

The identification of putative or novel ncRNAs in the mitochondria of human tissue and whether novel or known ncRNAs are differentially expressed in response to a physiological stressor such as endurance exercise has been an unexplored field of research. Exercise is the most potent non-pharmacological way to stimulate increased mitochondrial content in highly metabolic tissues such as skeletal muscle and heart. However, much of what is currently known about how mitochondria are regulated by stressors such as exercise is derived from measuring changes at the whole-cell level or is focussed on subcellular compartments other than the mitochondria, such as the cytosol and nucleus. The response of ncRNAs at the sub-cellular level is, however, largely unknown and is limited to a handful of studies conducted under pathological conditions. For example, following a diabetic insult in rodents, miR-378 expression in whole-cardiac muscle was unchanged, but increased in intermyofibrillar mitochondria, suggesting that ncRNAs may be redistributed between mitochondrial subpopulations in pathological states (19). This finding suggests that independent of whole cell ncRNA populations, mitochondrial ncRNAs may have advanced regulatory roles in both mitochondrial and whole cell metabolism. Therefore, a major finding of the present study was that differential expression analysis found that hsa-miR-146b-3p was significantly higher 3h post-exercise both in whole muscle and in isolated mitochondria. Hsa-miR-146b-3p displayed modest expression levels in the mitochondria 3h post-exercise but was detected at near-zero levels pre- and immediately post-exercise. In contrast, at the whole muscle level, hsa-miR-146b-3p was detected in all 36 samples (12 participants, 3 timepoints) and was differentially expressed in response to acute endurance exercise (Figure 4F). One explanation for these observations is that miRNAs may be transported in and out of the mitochondria in the early recovery phase following acute endurance exercise. This suggests that miRNAs may be targeted to the mitochondria from other parts of skeletal muscle cell in response to endurance exercise. At this stage, RNA and in particular miRNA transport across the mitochondrial membrane remains contentious. While it is generally accepted that miRNAs can co-localise in numerous subcellular comportments such as the nucleus, nucleolus, cytosol, and mitochondria (11, 44), the putative mechanisms illustrating how miRNAs are transported within cells remain elusive and have been discussed elsewhere (44).

With regards to miR-146b-3p, in silico analysis predicts 4091 nuclear encoded transcripts as targets based on sequence complementarity (TargetScanHuman 8.0 (45)). Although in silico analysis using miRwalk (46) returned 73 targets for miR-146b-3p that were experimentally validated using HITS-CLIP-seq, PAR-CLIP-seq, western blotting or luciferase reporter assays. there were no experimentally validated targets encoded by the mitochondrial genome. MiR-146b-3p has previously been reported to increase during both C2C12 myoblast differentiation (alongside the downregulation of *Smad4, Hmga2* and *Notch1*), and 3-5 days following muscle injury induced by the injection of barium chloride in the tibialis anterior of 10-week-old mice (47). These authors further confirmed direct interactions between miR-146b and the miRNA response elements in the 3’ UTR of *Smad4, Hmga2* and *Notch1*, identifying miR-146 as a regulator of myoblast differentiation and muscle regeneration (47). However, these experiments were conducted at the whole tissue level and the regulation of mitochondria-specific functions by miR-146b in skeletal muscle tissue have not yet been reported. More recently, miR-146b was shown to directly target mitogen-activated protein kinase kinase kinase 10 (MAP3K10) and promote cancer development following its downregulation in pancreatic cancer tissues (48). While cancer pathophysiology includes a shift towards glycolysis that is maintained under aerobic conditions and ultimately leads to mitochondrial dysfunction (49, 50), further research, such as via AGO-CLIP-based sequencing, should next seek to confirm potential interactions between miR-146b-3p and mitochondrial-encoded transcripts in skeletal muscle. Gain- or loss-of-function experiments in animal skeletal muscle tissue or cells are similarly required to further detail the physiological relevance of miR-146b-3p as it relates to mitochondria function in healthy tissue.

MiR-1 is a well characterised miRNA that is highly abundant in skeletal muscle tissue (43) and is crucial for skeletal muscle development, maintenance, remodelling, and the regulation of mitochondrial function (11, 51). The present study detected hsa-miR-1-3p in whole skeletal muscle at levels approximately 25-fold higher than any other miRNA. Hsa-miR-1-3p was also the most highly expressed species in mitochondria isolated from human skeletal muscle and was detected at levels approximately 7-fold higher than any other miRNA. This is consistent with earlier findings that miR-1, which is conserved between mice and humans, was particularly enriched in mitochondria isolated from C2C12 myotubes (20). Evidence from the current study corroborates previous observations that miR-1 can be detected in the mitochondria of human muscle precursor cells (28), and now in mitochondria isolated from human skeletal muscle tissue. Zhang and colleagues (20) employed crosslinking immunoprecipitation coupled with deep sequencing to demonstrate direct interactions of miR-1 with AGO2, a crucial component of RNA-induced silencing complexes, in the mitochondria of mouse muscle cells. Although miRNAs usually suppress protein translation, these authors found that miR-1 instead increased transcript and subsequently protein abundance for *Mt-Co1* and *Mt-Nd1* in C2C12 myocytes during differentiation and that AGO2, but not GW182, was required for enhanced mitochondria translation (20). In contrast, inhibition of miR-1 reduced *ND1* and *CO1* translation in C2C12 myoblasts in vitro (20). However, we did not observe changes in either direction for miR-1-3p, *Mt-Nd1* and *Mt-Co1* at the chosen time-points in the human acute exercise model described here. While direct interactions are yet to be demonstrated between AGO2, miR-1-3p and any putative targets of miR-1-3p in human mitochondria, the abundance of miR-1-3p in human skeletal muscle mitochondria as well as the previously documented interactions with mitochondrial-encoded transcripts (20) suggest that miR-1-3p may play a pivotal role in the key structural and regulatory processes in human skeletal muscle despite of lack of regulation in the early adaptive response to endurance training.

There are well described technical challenges in obtaining pure mitochondrial RNA. A cell fractionation approach in combination with RNase digestion is commonly used but can result in false positives due to incomplete RNase digestion (3, 52). To counter this, a refinement of the fractionation approach is to first remove the outer mitochondrial membrane and create mitoplasts prior to the RNase treatment, thus removing a significant source of non-mitochondrial RNA (3). In a series of experiments, we have recently established that mitochondria isolated via immunoprecipitation followed by RNase-A and proteinase-K treatments yields a transcriptome of similar purity to that of mitoplasts (32). Utilising this approach in the present study provides high confidence that only RNA species contained within the mitochondrial membranes have been identified.

It is currently contentious if several of the well described myo-miRs display differential expression in whole skeletal muscle following acute endurance exercise. We and others have previously reported expression levels of the muscle-enriched miR-1, miR-133a, miR-133b and miR-181a are increased in humans (15, 16), after one hour of moderate intensity cycling. However, we have also observed no increase in miR-1, miR-133a and miR-133b expression following comparable exercise protocols (53). All these aforementioned studies have measured expression using qPCR and utilised accepted approaches at the time (54), with RNU48 and/or U6 as endogenous controls because they were stably expressed. Those studies that did observe the expression of these miRs to significantly increase following acute exercise typically relied on normalising to a single endogenous control (15, 16) whereas the study that observed no increase in miR-1, miR-133a and miR-133b expression normalised to the median of two endogenous controls (53). RNA sequencing offers a more robust approach to normalisation as it calculates the geometric mean for each gene across all samples and internally corrects for library size prior to differential expression analysis (36). In the absence of RNA sequencing being available, qPCR approaches for quantifying miRNA expression should normalise to the median of two or more endogenous controls.

In summary, this study reports that mitochondria isolated from healthy male skeletal muscle contains a distinct population of approximately 110 miRNAs with several of the canonical myo-miRs comprising the bulk of the total mitochondrial miRNA reads. This study also presents evidence that the expression of miR-146b-3p is significantly increased in skeletal muscle whole tissue and mitochondria in the hours following a bout of endurance exercise suggesting that it may play a role in the regulation of exercise-induced mitochondrial adaptations. Future research is now required to investigate miRNA-mRNA interactions in the mitochondria of skeletal muscle tissue to assess the physiological relevance of mitochondria-localised miRNAs in response to positive metabolic stressors such as endurance exercise.

## Data Availability

RNA-sequencing datasets from this study have been deposited at the NCBI Gene Expression Omnibus for the whole-transcriptome in whole skeletal muscle (GSE276889) and small RNA sequencing for whole skeletal muscle (GSE277549) and for isolated mitochondria (GSE220280).

## Competing interests statement

The authors have no competing interests in connection with this article.

## Author Contributions

Conceptualisation: JLS, GDW, S.Lamon

Participant Recruitment and Exercise Testing: JLS

Data Collection: JLS, GDW

Data analysis: JLS, S.Loke

Data interpretation: JLS, LC, DH, MS

Writing original draft: JLS, S.Lamon, GDW

Review and approve the final manuscript: all authors.

## Acknowledgements

We acknowledge Dr Adam Trewin, Mr Pratik Churi and Ms Ashwinder Goshel for their assistance during participant testing. We acknowledge Dr Andrew Garnham (MD) for his involvement during the collection of skeletal muscle biopsies. Severine Lamon is supported by an Australian Research Council Future Fellowship (FT10100278).

## Supporting Information

**Supporting Information Figure 1.**
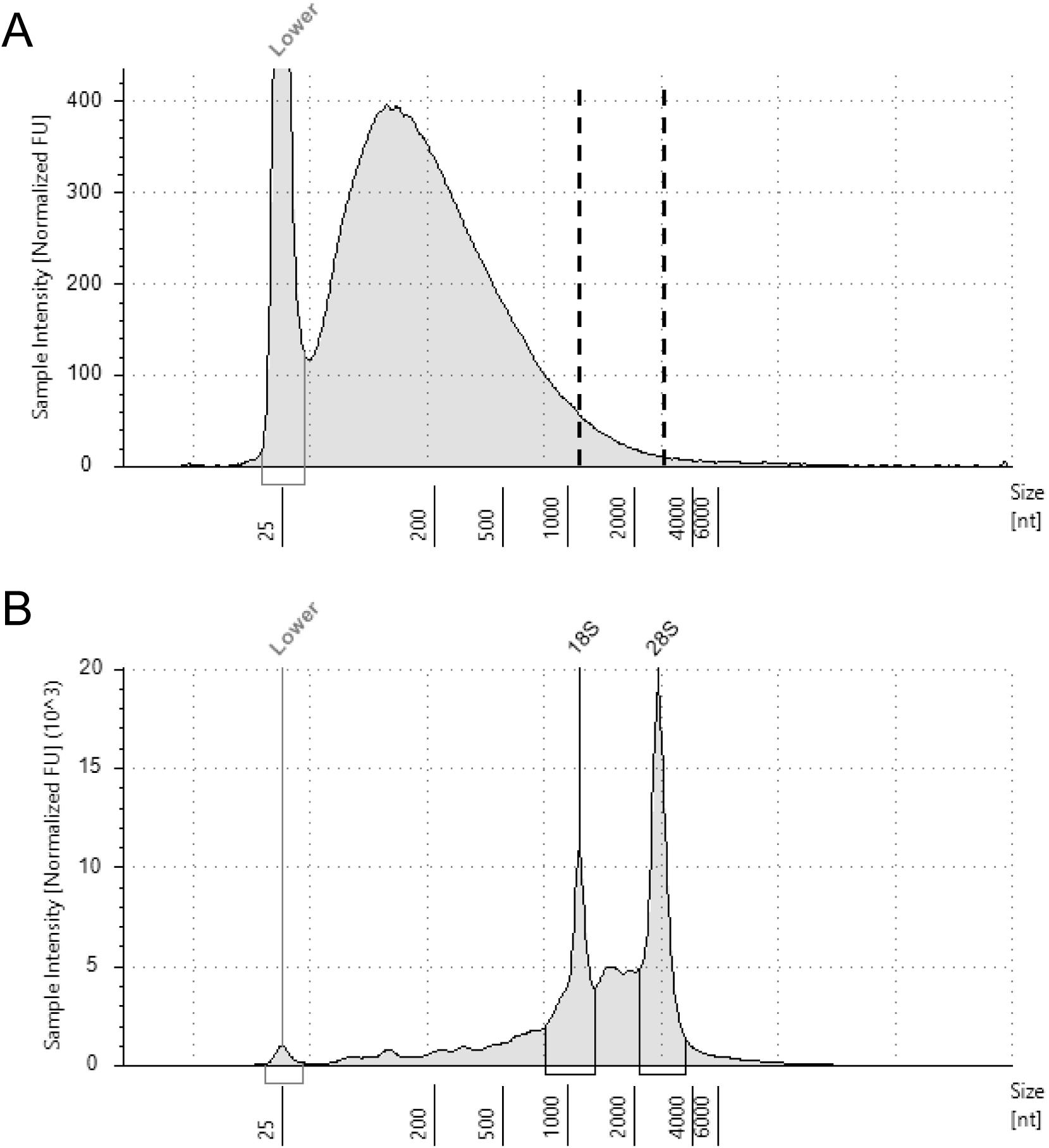
Representative electropherogram (cat# 5067-5579 and 5067-5580, HS RNA screentape, Agilent Technologies) of total RNA extracted from A) human skeletal muscle mitochondria and B) human whole skeletal muscle tissue. An absence of peaks at approximately 1000 and 3000 nt (dashed lines corresponding to 18s and 28s rRNA, respectively) suggests the mitochondrial RNA is free of non-mitochondrial RNAs. Peak at 25nt corresponds to an internal size standard.

**Supporting Information Figure 2.**
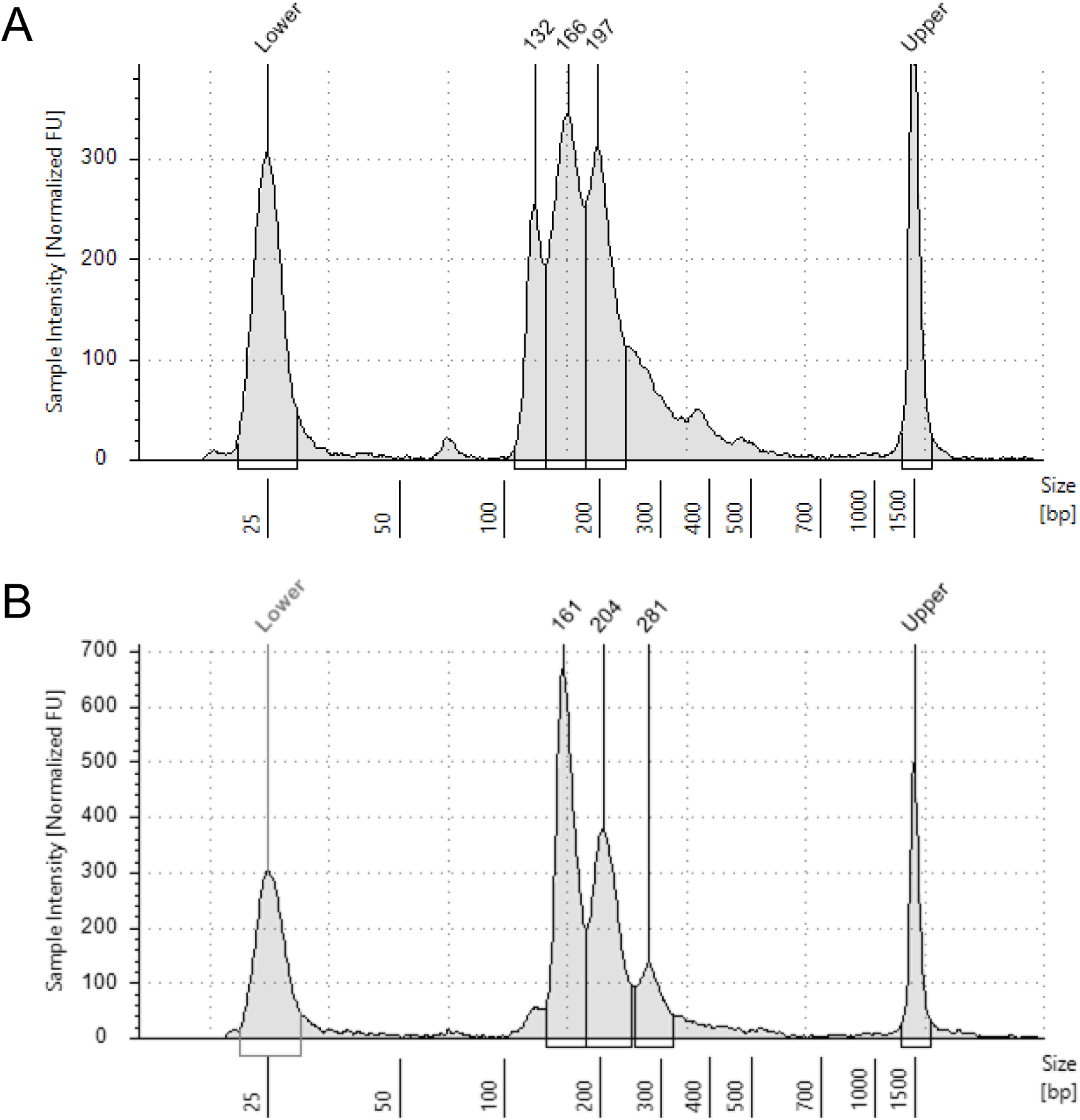
Representative electropherogram (HS D1000, cat# 5067-5584 and 5067-5585, Agilent Technologies) of small RNA libraries prepared from total mitochondrial RNA extracted from human skeletal muscle tissue. Adapter-ligated miRNAs are observed at approximately 160bp, while adapter-dimer and other small RNAs are observed at approximately 130 and 200-300bp, respectively. Peaks at 25 and 1500bp correspond to internal size standards.

**Supporting Information Figure 3.**
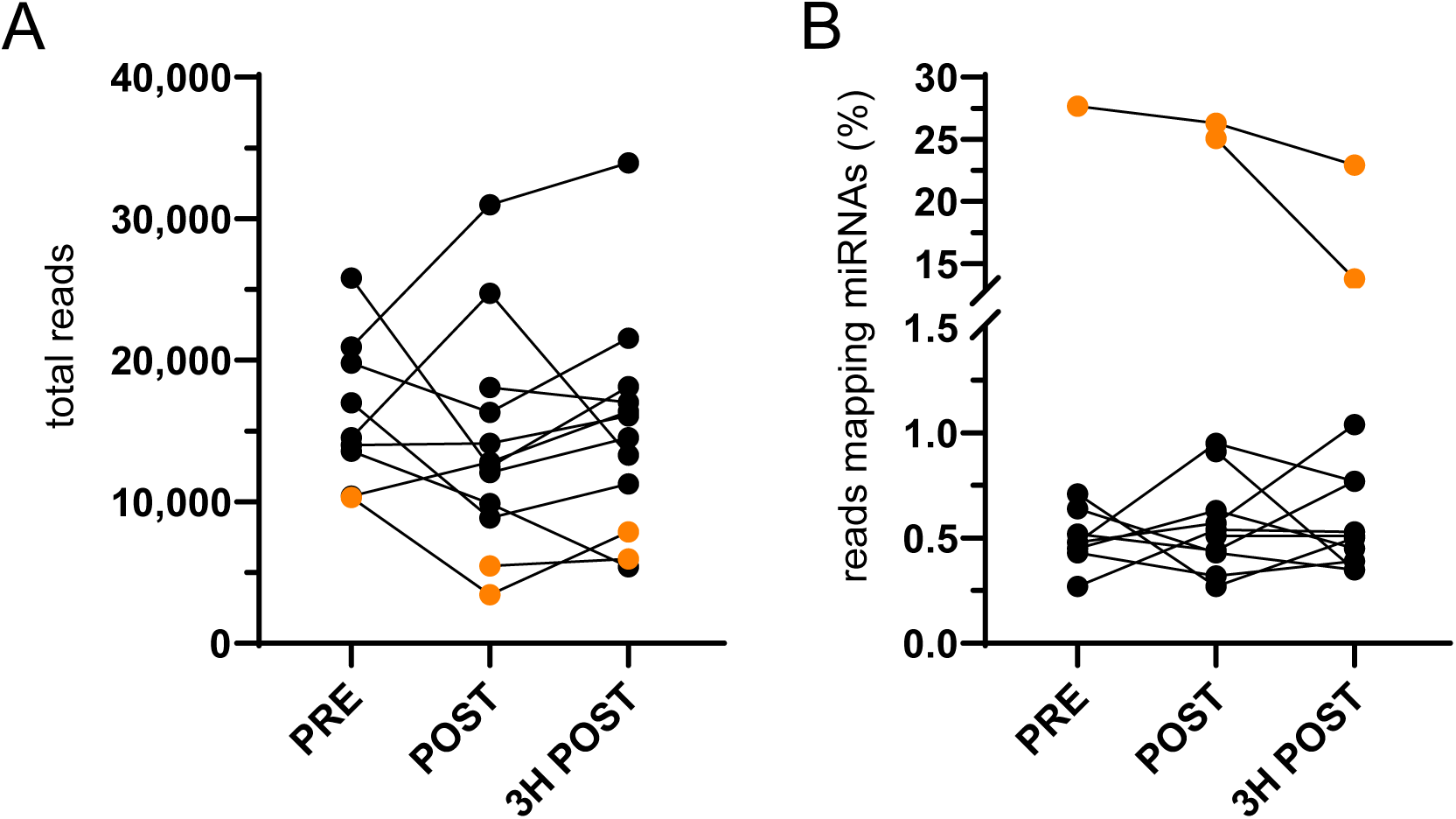
Small RNA libraries prepared from skeletal muscle mitochondria pre-, post- or 3h post-exercise and were pre-sequenced on a MiSeq Reagent Nano Kit v2 (MS-102-2001 Illumina, SD, USA) single-end 1×51 bp run. A) The total number of sequenced reads (p=0.69), and B) proportion of reads mapping mature human miRNAs were not significantly different between libraries. For five small RNA libraries highlighted in orange, the proportion of reads mapping miRNAs was between 20-30% and was well above the range presented for most samples. One-way, repeated measures ANOVAs with p-value <0.05 were used to identify significant differences.

**Supporting Information - Table 1.** Analysis of differential miRNA expression in whole skeletal muscle tissue pre-, immediately post- and 3h post-exercise. Differential expression analysis conducted using DEseq2 (v1.36) in RStudio (v4.2.1). Showing all miRNAs that returned a raw p-value of less than 0.05 prior to FDR adjustment. p (raw), raw p-value; FDR, false discovery rate or adjusted p-value (Benjamini and Hochberg correction for multiple testing); log2FC, log2 fold-change.

| post- vs pre-exercise |  |  |  |  | 3h post- vs pre-exercise |  |  |  |  | 3h post- vs post-exercise |  |  |  |  |
| --- | --- | --- | --- | --- | --- | --- | --- | --- | --- | --- | --- | --- | --- | --- |
|  | miRNA | p (raw) | FDR | log2FC |  | miRNA | p (raw) | FDR | log2FC |  | miRNA | p (raw) | FDR | log2FC |
| Up | hsa-miR-223-3p | <0.001 | <0.001 | 3.50 | Up | hsa-miR-223-3p | <0.001 | <0.001 | 3.84 | Up | hsa-miR-146b-3p | <0.001 | 0.030 | 2.33 |
|  | hsa-miR-223-5p | <0.001 | 0.004 | 2.51 |  | hsa-miR-223-5p | <0.001 | <0.001 | 3.17 |  | hsa-miR-221-5p | 0.001 | 0.228 | 1.08 |
|  | hsa-miR-15b-5p | <0.001 | 0.030 | 1.12 |  | hsa-miR-146b-3p | <0.001 | 0.072 | 1.97 |  | hsa-miR-146b-5p | 0.020 | 0.986 | 0.79 |
|  | hsa-miR-106b-3p | <0.001 | 0.040 | 0.73 |  | hsa-miR-142-5p | 0.005 | 0.435 | 1.12 |  | hsa-miR-4676-3p | 0.043 | 0.986 | 1.20 |
|  | hsa-miR-142-5p | 0.004 | 0.302 | 1.15 |  | hsa-miR-324-3p | 0.013 | 0.847 | 0.88 |  | hsa-miR-92a-1-5p | 0.043 | 0.986 | 1.23 |
|  | hsa-miR-3615 | 0.004 | 0.309 | 0.68 |  | hsa-miR-338-5p | 0.017 | 0.902 | 0.82 |  | hsa-miR-203a-3p | 0.043 | 0.986 | 1.79 |
|  | hsa-miR-454-3p | 0.009 | 0.353 | 0.72 |  | hsa-miR-3065-3p | 0.019 | 0.902 | 0.80 |  | hsa-miR-203b-5p | 0.043 | 0.986 | 1.79 |
|  | hsa-miR-584-5p | 0.009 | 0.353 | 0.51 |  | hsa-miR-3615 | 0.019 | 0.902 | 0.55 |  | hsa-miR-1247-3p | 0.045 | 0.986 | 1.73 |
|  | hsa-miR-652-3p | 0.013 | 0.399 | 0.50 |  | hsa-miR-92a-1-5p | 0.023 | 0.920 | 1.42 |  | hsa-miR-335-3p | 0.050 | 0.986 | 0.40 |
|  | hsa-miR-503-5p | 0.016 | 0.400 | 0.87 |  | hsa-miR-503-5p | 0.024 | 0.920 | 0.82 |  |  |  |  |  |
|  | hsa-miR-618 | 0.016 | 0.400 | 1.64 |  | hsa-miR-7976 | 0.024 | 0.920 | 1.80 |  |  |  |  |  |
|  | hsa-miR-941 | 0.017 | 0.400 | 0.54 |  | hsa-miR-185-5p | 0.027 | 0.920 | 0.52 |  |  |  |  |  |
|  | hsa-miR-185-3p | 0.017 | 0.406 | 0.74 |  | hsa-miR-6894-3p | 0.028 | 0.920 | 1.66 |  |  |  |  |  |
|  | hsa-miR-16-5p | 0.019 | 0.414 | 0.59 |  | hsa-miR-20b-5p | 0.033 | 0.920 | 0.69 |  |  |  |  |  |
|  | hsa-miR-1-5p | 0.019 | 0.414 | 0.94 |  | hsa-miR-106b-3p | 0.035 | 0.920 | 0.42 |  |  |  |  |  |
|  | hsa-miR-550a-5p | 0.021 | 0.414 | 1.47 |  | hsa-miR-93-5p | 0.038 | 0.920 | 0.55 |  |  |  |  |  |
|  | hsa-miR-93-5p | 0.022 | 0.414 | 0.61 |  | hsa-miR-103a-3p | 0.041 | 0.920 | 0.35 |  |  |  |  |  |
|  | hsa-miR-92a-3p | 0.024 | 0.414 | 0.46 |  | hsa-miR-103b | 0.041 | 0.920 | 0.35 |  |  |  |  |  |
|  | hsa-miR-451b | 0.026 | 0.414 | 1.39 |  | hsa-miR-107 | 0.042 | 0.920 | 0.35 |  |  |  |  |  |
|  | hsa-miR-25-3p | 0.034 | 0.482 | 0.41 |  | hsa-miR-17-5p | 0.043 | 0.920 | 0.56 |  |  |  |  |  |
|  | hsa-miR-451a | 0.034 | 0.482 | 0.75 |  | hsa-miR-106a-5p | 0.045 | 0.920 | 0.56 |  |  |  |  |  |
|  | hsa-miR-589-5p | 0.036 | 0.482 | 0.50 |  | hsa-miR-15b-5p | 0.045 | 0.920 | 0.60 |  |  |  |  |  |
|  | hsa-miR-550a-3-5p | 0.037 | 0.482 | 1.31 |  | hsa-miR-191-5p | 0.045 | 0.920 | 0.30 |  |  |  |  |  |
|  | hsa-miR-324-3p | 0.042 | 0.521 | 0.73 |  | hsa-miR-221-5p | 0.047 | 0.923 | 0.61 |  |  |  |  |  |
|  | hsa-miR-144-5p | 0.045 | 0.528 | 0.80 |  |  |  |  |  |  |  |  |  |  |
|  | hsa-miR-20b-5p | 0.047 | 0.528 | 0.64 |  |  |  |  |  |  |  |  |  |  |
|  | hsa-miR-4521 | 0.048 | 0.528 | 0.69 |  |  |  |  |  |  |  |  |  |  |
| Down | hsa-miR-452-5p | 0.000 | 0.030 | -1.49 | Down | hsa-miR-340-3p | <0.001 | 0.072 | -0.95 | Down | hsa-miR-550a-5p | 0.016 | 0.986 | -1.55 |
|  | hsa-miR-139-5p | 0.001 | 0.097 | -0.52 |  | hsa-miR-139-5p | <0.001 | 0.072 | -0.54 |  | hsa-miR-550a-3-5p | 0.021 | 0.986 | -1.47 |
|  | hsa-let-7a-3p | 0.001 | 0.161 | -0.84 |  | hsa-miR-654-3p | 0.001 | 0.173 | -0.61 |  | hsa-miR-550b-3p | 0.029 | 0.986 | -1.40 |
|  | hsa-miR-208b-3p | 0.003 | 0.264 | -0.99 |  | hsa-miR-148a-3p | 0.002 | 0.246 | -0.70 |  | hsa-miR-323a-3p | 0.045 | 0.986 | -0.53 |
|  | hsa-miR-374b-3p | 0.005 | 0.353 | -0.99 |  | hsa-miR-3909 | 0.009 | 0.681 | -0.46 |  | hsa-miR-425-5p | 0.045 | 0.986 | -0.51 |
|  | hsa-miR-374a-3p | 0.006 | 0.353 | -1.17 |  | hsa-miR-708-3p | 0.010 | 0.681 | -1.17 |  |  |  |  |  |
|  | hsa-miR-148a-3p | 0.007 | 0.353 | -0.62 |  | hsa-let-7a-3p | 0.010 | 0.681 | -0.68 |  |  |  |  |  |
|  | hsa-miR-382-3p | 0.007 | 0.353 | -0.87 |  | hsa-miR-452-5p | 0.016 | 0.902 | -0.92 |  |  |  |  |  |
|  | hsa-miR-335-3p | 0.007 | 0.353 | -0.55 |  | hsa-let-7f-2-3p | 0.020 | 0.902 | -1.99 |  |  |  |  |  |
|  | hsa-miR-199a-3p | 0.009 | 0.353 | -0.53 |  | hsa-miR-374a-3p | 0.032 | 0.920 | -0.91 |  |  |  |  |  |
|  | hsa-miR-199b-3p | 0.009 | 0.353 | -0.53 |  | hsa-miR-126-3p | 0.037 | 0.920 | -0.30 |  |  |  |  |  |
|  | hsa-miR-34c-5p | 0.009 | 0.353 | -1.51 |  | hsa-miR-128-3p | 0.041 | 0.920 | -0.38 |  |  |  |  |  |
|  | hsa-miR-29c-3p | 0.011 | 0.399 | -0.85 |  | hsa-miR-323a-3p | 0.042 | 0.920 | -0.54 |  |  |  |  |  |
|  | hsa-miR-660-5p | 0.013 | 0.399 | -0.66 |  | hsa-miR-27a-3p | 0.043 | 0.920 | -0.37 |  |  |  |  |  |
|  | hsa-miR-126-3p | 0.013 | 0.399 | -0.35 |  | hsa-miR-1285-5p | 0.045 | 0.920 | -1.83 |  |  |  |  |  |
|  | hsa-miR-654-3p | 0.013 | 0.399 | -0.47 |  |  |  |  |  |  |  |  |  |  |
|  | hsa-miR-203a-3p | 0.015 | 0.400 | -2.17 |  |  |  |  |  |  |  |  |  |  |
|  | hsa-miR-203b-5p | 0.015 | 0.400 | -2.17 |  |  |  |  |  |  |  |  |  |  |
|  | hsa-miR-24-2-5p | 0.015 | 0.400 | -0.62 |  |  |  |  |  |  |  |  |  |  |
|  | hsa-miR-769-3p | 0.016 | 0.400 | -2.20 |  |  |  |  |  |  |  |  |  |  |
|  | hsa-miR-340-5p | 0.019 | 0.414 | -0.57 |  |  |  |  |  |  |  |  |  |  |
|  | hsa-miR-499a-5p | 0.021 | 0.414 | -0.64 |  |  |  |  |  |  |  |  |  |  |
|  | hsa-miR-499b-3p | 0.022 | 0.414 | -0.64 |  |  |  |  |  |  |  |  |  |  |
|  | hsa-let-7f-2-3p | 0.022 | 0.414 | -1.96 |  |  |  |  |  |  |  |  |  |  |
|  | hsa-miR-4796-5p | 0.023 | 0.414 | -2.34 |  |  |  |  |  |  |  |  |  |  |
|  | hsa-miR-27a-3p | 0.024 | 0.414 | -0.41 |  |  |  |  |  |  |  |  |  |  |
|  | hsa-miR-9-5p | 0.025 | 0.414 | -0.84 |  |  |  |  |  |  |  |  |  |  |
|  | hsa-miR-3605-5p | 0.025 | 0.414 | -1.31 |  |  |  |  |  |  |  |  |  |  |
|  | hsa-miR-21-5p | 0.025 | 0.414 | -0.39 |  |  |  |  |  |  |  |  |  |  |
|  | hsa-miR-1292-5p | 0.028 | 0.441 | -1.46 |  |  |  |  |  |  |  |  |  |  |
|  | hsa-miR-199b-5p | 0.031 | 0.482 | -0.53 |  |  |  |  |  |  |  |  |  |  |
|  | hsa-miR-340-3p | 0.032 | 0.482 | -0.56 |  |  |  |  |  |  |  |  |  |  |
|  | hsa-miR-99a-5p | 0.033 | 0.482 | -0.36 |  |  |  |  |  |  |  |  |  |  |
|  | hsa-miR-30a-5p | 0.034 | 0.482 | -0.36 |  |  |  |  |  |  |  |  |  |  |
|  | hsa-miR-455-5p | 0.035 | 0.482 | -0.38 |  |  |  |  |  |  |  |  |  |  |
|  | hsa-miR-4662b | 0.036 | 0.482 | -0.96 |  |  |  |  |  |  |  |  |  |  |
|  | hsa-miR-4662a-5p | 0.037 | 0.482 | -0.96 |  |  |  |  |  |  |  |  |  |  |
|  | hsa-miR-10395-3p | 0.040 | 0.508 | -0.59 |  |  |  |  |  |  |  |  |  |  |
|  | hsa-miR-337-3p | 0.042 | 0.521 | -1.40 |  |  |  |  |  |  |  |  |  |  |
|  | hsa-miR-493-3p | 0.046 | 0.528 | -0.60 |  |  |  |  |  |  |  |  |  |  |
|  | hsa-miR-335-5p | 0.046 | 0.528 | -1.35 |  |  |  |  |  |  |  |  |  |  |
|  | hsa-miR-145-3p | 0.047 | 0.528 | -0.50 |  |  |  |  |  |  |  |  |  |  |
|  | hsa-miR-944 | 0.048 | 0.528 | -0.45 |  |  |  |  |  |  |  |  |  |  |

**Supporting Information - Table 2.**
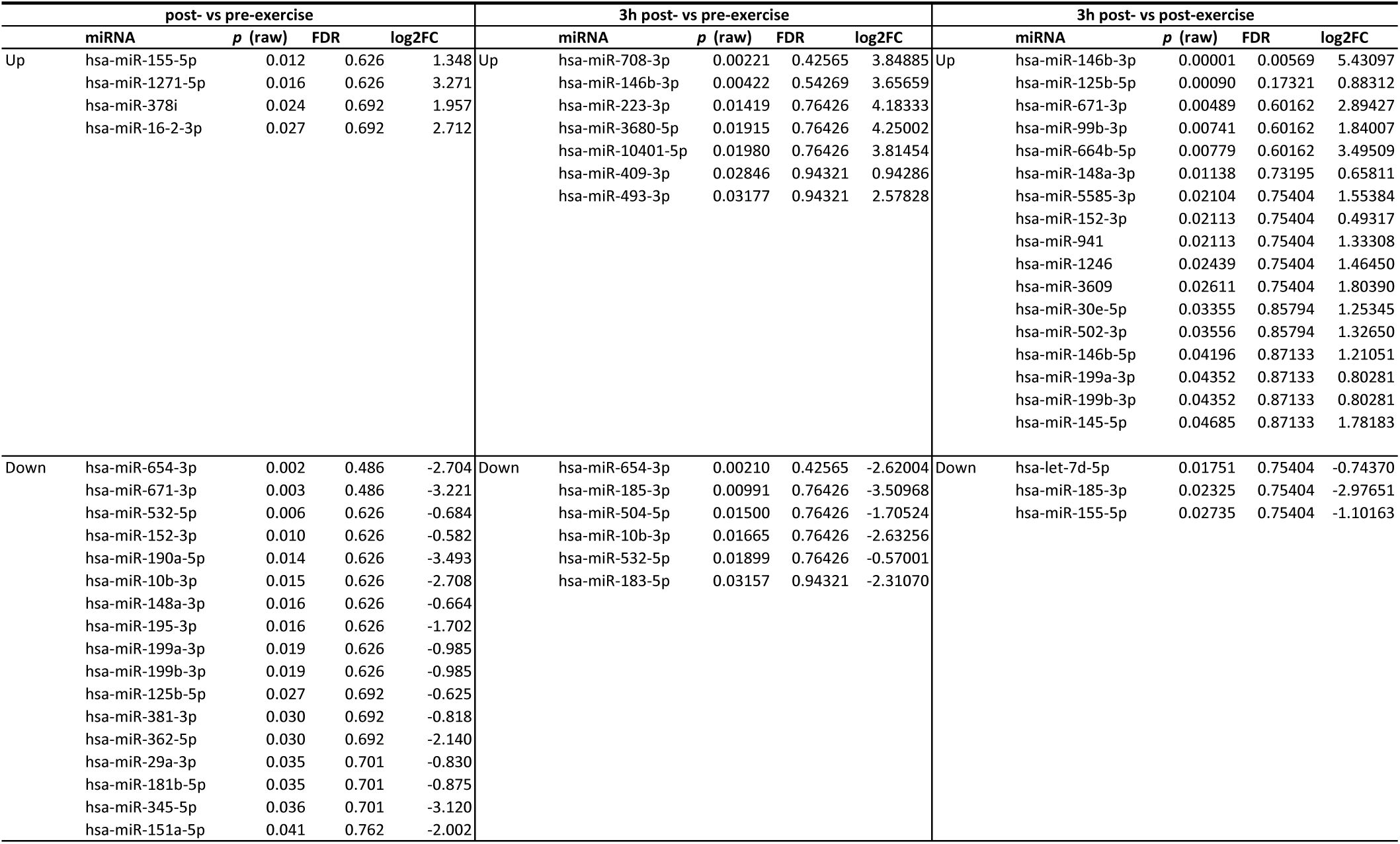
Analysis of differential miRNA expression in skeletal muscle mitochondria pre-, immediately post- and 3h post-exercise. Differential expression analysis conducted using DEseq2 (v1.36) in RStudio (v4.2.1). Showing all miRNAs that returned a raw p-value of less than 0.05 prior to FDR adjustment. p (raw), raw p-value; FDR, false discovery rate or adjusted p-value (Benjamini and Hochberg correction for multiple testing); log2FC, log2 fold-change.

## Notes

### Competing Interest Statement

The authors have declared no competing interest.

